# Accessing rare bacterial biosphere of soil through culturing: a comparative study of culture media effectiveness integrated with metataxonomics

**DOI:** 10.1101/2025.10.20.683382

**Authors:** José A. Siles, Norman Terry

**Author notes:** Department of Soil and Water Conservation and Organic Waste Management, Centro de Edafología y Biología Aplicada del Segura-Consejo Superior de Investigaciones Científicas, CEBAS-CSIC, 30100 Murcia, Spain.

## Abstract

Soil harbors a vast bacterial diversity, including a “rare biosphere” of low-abundance taxa increasingly recognized as crucial for ecosystem functioning. Traditional plate-based culturing can facilitate access to this fraction. Here, we investigated how culture medium type influences the recovery of rare soil bacteria. To do this, culture-independent bacterial diversity of agricultural, forest, and contaminated soils was compared with culture-dependent diversity on four media (TSA (tryptic soy agar), SEM (soil extract medium), R2A (Reasoner’s 2A), and 20-fold diluted R2A) using a high-throughput approach in which all colonies were collected and analyzed via 16S rRNA gene metabarcoding (metataxonomics). Across all media and soils, ~30% of the culture-independent taxa were recovered through culturing, of which ~40% were members of the rare community. Culturability of bacterial communities declined across soil types in the order: contaminated > agricultural > forest, but the community isolated from forest soil contained a higher proportion of rare taxa. SEM, characterized by its low nutrient content and chemical composition resembling that of soil, proved to be the most effective, yielding higher colony-forming unit counts, greater overall and rare richness, and recovering a more taxonomically diverse rare community. The rare community retrieved by SEM was dominated by non–spore-forming taxa, in contrast to the other media, including diluted R2A. Culturing revealed a unique fraction of culturable taxa, representing ~17% of the culture-independent community on average, likely members of the rare biosphere that escape detection in metabarcoding analyses. SEM was also the medium that best captured this fraction. Our study demonstrates that low-nutrient, soil-like media such as SEM are powerful tools for accessing and characterizing the rare soil biosphere, complementing metabarcoding and providing isolates to explore the ecological and functional significance of rare taxa.

## 1 Introduction

Soil has been estimated to harbor up to 59 % of the total known species of bacteria on Earth (Anthony et al., 2023). The investigation of this diversity is not only interesting in itself, but also because soil bacteria mediate processes that are key for ecosystem services, such as nutrient cycling, land productivity, contaminant degradation, pathogen control, and climate regulation (Delgado-Baquerizo et al., 2016; Köninger et al., 2022). The introduction of high-throughput sequencing approaches such as metataxonomics (also known as 16S rRNA gene metabarcoding) has greatly advanced our understanding and discovery of this diversity (Thompson et al., 2017). Additionally, soil 16S rRNA gene metabarcoding surveys typically reveal a long tail of low-relative-abundance taxa, which constitute the so-called bacterial rare biosphere (Lynch and Neufeld, 2015). In the last few years, the interest in this microbial fraction has increased since rare microbial taxa are now believed to be more relevant to ecosystem functioning than previously anticipated (Chen et al., 2020). First, because many rare taxa may respond to disturbances and can become abundant under changing conditions (Riddley et al., 2025). Second, the vast diversity of low-abundance taxa serves as a large reservoir of genetic traits that support a wide range of both established and potentially novel microbial functions. Third, members of the rare biosphere can exhibit disproportionately high activity relative to their abundance in performing key functions (Pascoal et al., 2021). In this context, for example, Pulido-Chavez et al. (2023) showed that rare bacteria taxa belonging to the genera *Massilia* and *Noviherbaspirillium* in the undisturbed soil became dominant in a chaparral after a wildfire due to their ability to exploit post-fire resources. Additionally, the growing interest in rare soil bacteria is largely driven by their potential in bioprospecting, particularly for developing novel biotechnological solutions (Pascoal et al., 2021; Pascoal et al., 2020). Metataxonomics identifies the species present at a given time and provides valuable insights into their potential ecological roles. Nonetheless, these predicted roles require experimental validation. In this regard, culture-dependent approaches remain essential for uncovering the ecological functions of rare soil bacteria (VanInsberghe et al., 2013).

In recent years, in concomitance with the growing focus on rare soil bacteria and the accumulation of sequencing information, a “renaissance” in culturing approaches has emerged (Carini, 2019; Thrash, 2021). This renewed motivation to culture bacteria from multiple ecosystems surges, in part, from our limited ability to annotate genes from bacterial genomes or metagenomes, an issue that can be most effectively addressed by isolating new bacterial strains and investigating their physiology and ecology (Carini, 2019; Thrash, 2019). This renewed interest in culturing is also driven by the successful isolation of novel bacterial taxa, which have attracted considerable attention either due to their unique characteristics or because they have advanced our understanding of key natural processes (Lewis et al., 2020). These advances have been made possible, in part, by the development of innovative culturing methods designed to recover bacteria that were previously considered unculturable (Lewis et al., 2020; Thrash, 2021). Despite these new methods, plate culturing with subsequent colony picking and identification remains the gold standard in most laboratories for culture-dependent studies of soil bacteria. Furthermore, the simplest way to maintain or handle newly isolated bacteria, including those obtained through novel culturing techniques, is by using solid-media culture plates or related devices. This highlights the ongoing need for research focused on plate-based culturing and the identification of media and conditions that support a higher bacterial diversity (Nguyen et al., 2018). Although the classical “1% culturability paradigm” is no longer accurate, most bacterial taxa remain unculturable using plate culturing (Molina-Menor et al., 2021; Steen et al., 2019). Several strategies can facilitate the isolation of novel bacterial strains using plate-based methods, including: (i) the use of media with contrasting nutrient concentrations, or supplemented with specific nutrients or chemical factors for bacterial growth; (ii) coculture with helper bacteria; (iii) extended incubation periods; (iv) alternating incubation temperatures; and (v) techniques to physically reduce the number and diversity of bacteria in the sample (Molina-Menor et al., 2021; Nguyen et al., 2018; Pham and Kim, 2012). This renewed interest in cultivating soil bacteria presents an opportunity to uncover the hidden diversity of rare soil taxa through improved culture-dependent, plate-based approaches, a topic that has not been adequately addressed in previous studies.

Several studies have evaluated the extent to which traditional culturing methods can access rare soil bacterial communities. To do this, 16S rRNA gene metabarcoding is combined with plate culturing and subsequent identification of isolates. Shade et al. (2012) and VanInsberghe et al. (2013) demonstrated that the accessibility of the rare bacteria biosphere via culturing is higher than initially anticipated. These studies employed various growth media for bacterial isolation but did not specifically assess how media type (in terms of nutrient contents) influences richness and assemblage of the isolated rare communities. However, the nutritional composition and content of culture media play a critical role in the isolation of rare bacteria, as they directly determine which organisms are able to grow (Bartelme et al., 2020). Rare taxa are thought to have specialized metabolic requirements that differ from those of abundant, fast-growing species, which tend to proliferate in nutrient-rich media (Kurm et al., 2019b). In addition, the rapid proliferation of these fast-growing species can outcompete and inhibit the growth of rare bacteria. Oligotrophic media in combination with long incubation periods are expected to increase the cultivability of rare bacteria. Extended incubation periods may be required for the reactivation of dormant rare species (Aanderud et al., 2015), as their delayed awakening can hinder their cultivability in short-term approaches. These expectations were not confirmed by Kurm et al. (2019a); nonetheless, this study was limited to a single soil type, biased toward small-celled bacteria, and based on only 172 isolates. Further studies are thus needed.

Some studies have found that when culture-dependent characterization of a soil bacterial community is combined with 16S rRNA gene metabarcoding, a substantial number of taxa are only detected by the culture-dependent approach (Lee et al., 2016; Mandakovic et al., 2018; Shade et al., 2012; Siles et al., 2022). Although methodological biases in 16S metabarcoding may contribute to this phenomenon, it has also been suggested that these taxa represent members of the rare bacterial community (Pédron et al., 2020). This unique culturable fraction of bacteria is hypothesized to exist in dormant or low-metabolic states, producing little DNA and thus evading detection by metabarcoding (Shade et al., 2012). Under suitable conditions, they can resume growth, allowing cultivation and revealing insights into the rare biosphere. For example, Siles et al. (2022) combined culturing and metataxonomics to study bacterial diversity in contaminated waste sediment and isolated a novel species from the rare biosphere with bioremediation potential, which was detected only through culturing. These findings highlight the complementary value of combining culture-independent and culture-dependent approaches in studying soil bacteria (Shade et al., 2012). Despite these important findings, previous studies have not thoroughly examined how richness and taxonomic composition of this culture-dependent fraction of potentially rare bacteria vary with culture media and across soil types.

This study aimed to examine how the type of culture medium influences the richness and taxonomic composition of rare soil bacteria isolated from contrasting soil ecosystems, with separate and detailed analyses of taxa both detected and undetected by culture-independent methods. To do this, we examined the culture-independent bacterial diversity of agricultural, forest, and contaminated soils and compared it with the culture-dependent diversity obtained using four media with different nutrient concentration and composition: TSA, SEM (soil-extracted medium), R2A, and 20-fold diluted R2A. In order to enable a direct comparison between culture-dependent and culture-independent communities, a high-throughput culturing approach was used, in which all bacterial isolates growing on the culture media were collected and their taxonomic composition analyzed through 16S rRNA gene metabarcoding. We hypothesized that SEM would recover a richer and more taxonomically diverse rare community in the three soils, as we expected this medium to be oligotrophic and to better reflect the nutrient content and composition required for soil bacteria to grow. We also anticipated that this medium will be the most effective at recovering unique culturable taxa absent from the culture-independent dataset, exhibiting a differentiated taxonomic composition and a higher proportion of unclassified sequences.

## 2 Materials and methods

### 2.1 Study sites, and soil sampling and characterization

Soil from three ecosystem types was considered in the present study: an agricultural field, a redwood forest, and a former multi-contaminated dumpsite. Agricultural soil was obtained from the Oxford Tract Field (37°52’34.6”N 122°16’01.7”W), a 2-ha research plot situated in Berkeley, California, and used by UC Berkeley staff to cultivate different types of seasonal crops (maize, sunflower, tomato, lettuce, etc.) for research purposes. At the time of sampling, the soil had been recently ploughed, and there were no plants present in the plot. The climate at the sampling site is typically Mediterranean, with a mean annual temperature of 14.6 °C and a mean annual precipitation of 737 mm.

Forest soil was sampled from a redwood forest in Tilden Regional Park, located in the Berkeley Hills of the San Francisco Bay Area, California. The park, with a size of 841 ha, is used for recreational purposes. Oak-Bay (*Quercus agrifoli* and *Umbellularia californica*) and Eucalyptus (*Eucalyptus globulus*) forests –with varying understory vegetation– predominate across the park, although areas of native coastal scrub are also present. Forest patches dominated by redwood (*Sequoia semperviren*s) can also be found in the park and one of them was selected for the present study (37°53’59.3”N 122°15’33.3”W). At the sampled forest site, understory vegetation was sparce due to the closed canopies and the soil presented an abundant litter layer, which was removed for soil sampling. The sampling site has a typical Mediterranean climate, characterized by a mean annual temperature of 14.6 °C and an average annual precipitation of 737 mm.

Contaminated soil was collected from a former industrial dumpsite located in Bay Point, California, USA (38°02’12.7”N, 121°56’52.9”W). This 29-ha former dumpsite was used to receive stormwater and wastewater discharges from commercial production plants. The site is contaminated with total petroleum hydrocarbons (20.14 g kg^−1^), polycyclic aromatic hydrocarbons (370 mg g^−1^), metals (chromium, barium, and zinc, among others) and carbon black (fly ash residue from coal burning industry, particle size <10 μm). The site is irregularly colonized by saltgrass (*Distichlis spicata*), pickleweed (*Salicornia pacifica*), and sea purslane (*Sesuvium verrucosum*). The sampling site experiences a Mediterranean-type climate, with an average annual temperature of 15.7 °C and an annual precipitation of 441 mm.

At each site, four independent replicate plots (5 × 5 m; about 100 m apart) were selected and one sample was collected from each plot. Each sample was a composite of five subsamples taken from the top 20 cm of soil: four subsamples collected orthogonally within a 2-m radius around a central subsample. In total, our study consisted of 12 replicate soil samples (3 soil types × 4 replicates). Then, samples were transported to the lab in cooled boxes and subsequently homogenized and sieved (2 mm mesh). Samples were separated into two portions: one was stored at 4 °C for physicochemical characterization and bacterial plate culturing, and the other was frozen (−80 °C) until DNA extraction.

Soil samples were physicochemically characterized (texture, pH, electrical conductivity, total and organic carbon (TC, TOC), total nitrogen (N), and phosphorus (P)) following standard methods at the UC Davis Analytical Lab (Davis, CA, USA). A summary of the results is provided in Table S1.

### 2.2 Culture-independent approach

For the assessing of the culture-independent bacterial diversity, total DNA from the 12 soil replicate samples was extracted in triplicate from 250 mg of soil mass using the Dneasy PowerSoil Kit (Qiagen) following the manufacturer’s instructions. Triplicate DNA extracts from each sample were then pooled and DNAs were purified using the DNA Clean & Concentrator Kit (Zymo Research) according to manufacturer’s instructions. Quality of DNAs was checked as previously described and DNA concentrations were normalized prior to 16S rRNA gene amplicon sequencing.

### 2.3 Culture-dependent approach

In each soil, culture-dependent bacterial diversity was investigated through plate culturing using four culture media differing in their nutrient concentrations: tryptic soy agar (TSA; BactoTM Tryptic Soy Broth. Soybean-Casein Digest Medium Becton, Dickinson and Company), Reasoner’s 2A (R2A, Research Products International) at standard concentration, R2A medium diluted 20-fold (1:20 R2A) and soil extract medium (SEM). SEM was prepared following a modified version of the method described in the DSMZ Media Catalogue (Leibniz Institute). Briefly, 400 g of air-dried soil were mixed with 1 L of distilled water, and the suspension was subsequently shaken for 12 hours. The suspension was then autoclaved for one hour at 121 °C, filtered, and supplemented with agar before sterilizing it at 121 °C for 20 minutes. For each soil, four media were prepared, with their pH adjusted to match that of the corresponding soil. Cycloheximide (200 μg ml^−1^) was used to exclude fungal growth. Contents of total organic C, N, P, potassium (K), calcium (Ca), magnesium (Mg), and sodium (Na) were determined in each culture media following standard methods at the UC Davis Analytical Lab (Davis, CA, USA).

For bacteria plate culturing, 4 g of each replicate soil sample were diluted with 36 mL of sterile saline solution (0.85% NaCl, w/v), vortexed for 1 min, sonicated for 15 seconds in a bath sonicator, and shaken for 90 minutes. Soil suspensions were then allowed to settle for 5 minutes, and dilutions were made with saline solutions. For each soil replicate, triplicate series of 100 µl of the 10^−4^, 10^−5^, and 10^−6^ dilutions were plated (3 soils × 4 replicate samples × 4 culture media × 3 dilutions × 3 replicates = 432 plates). These dilutions were selected based on preliminary tests to prevent overgrowth. The plates were sealed with Parafilm and incubated at alternating temperatures of 10 and 20 °C for 54 days. After this period, no new colony growth was observed, and incubation was discontinued. Visible colonies were counted and colony forming units (CFU) were calculated. Further, bacterial biomass growing on the plates was collected and homogenized with a spatula, and subsequently stored at −80 °C for DNA extraction. For each soil replicate sample and culture media, biomass of the different dilutions was merged. In total, the culture-dependent assay consisted of 48 samples (3 soils × 4 replicate samples × 4 culture media = 48 samples) prior DNA extraction with the kit *Quick*-DNA Fungal/Bacterial Kit (Zymo Research). DNAs were then purified using the DNA Clean & Concentrator Kit (Zymo Research). The quality of the DNAs was spectrophotometrically checked by NanoDrop (Thermo Fisher Scientific) based on the absorbance ratios A260/A280 and A260/A230. Subsequently, DNAs were quantified using QuantiFluor™ dsDNA System (Promega) and DNA concentrations were standardized prior to 16S rRNA gene amplicon sequencing.

### 2.4 16S rRNA gene amplicon sequencing and bioinformatic processing of the sequences

Bacterial culture-independent and -dependent diversity was investigated through 16S rRNA gene amplicon sequencing of 60 DNA samples. A fragment of the 16S rRNA gene, capturing the V1-V3 regions, was amplified using the primers 27F (5′-AGRGTTTGATCMTGGCTCA-3′) and 519R (5′-CCCCGYCAATTCMTTTRAGT-3′). PCR reactions were conducted using the HotStarTaq *Plus* Master Mix Kit (Qiagen) and barcoded forward primers, under the following thermal conditions: (i) 94°C for 3 min; (ii) 30 cycles of 94°C for 30 s, 53°C for 40 s, and 72°C for 60s; with (iii) a final elongation step at 72°C for 5 min. The success of the amplifications was checked in a 2% agarose gel and PCR products were subsequently purified using Agencourt AMPure XP magnetic beads kit (Beckman Coulter), quantified with the QuantiFluor™ dsDNA System (Promega), and pooled in equal proportions. The pooled product was then used to prepare the Illumina DNA library. Paired-end (PE) sequencing (2 × 300) was performed on an Illumina MiSeq sequencing platform at MR DNA (www.mrdnalab.com, Shallowater, TX, USA).

Bioinformatic processing of the sequences was conducted using the USEARCH pipeline and UPARSE-OTU algorithm (Edgar, 2013). Briefly, PE sequences were firstly merged with the command - *fastq_mergepairs*. Then, reads were quality-filtered allowing a maximum e-value of 1.0, trimmed to 450-bp (base pair), dereplicated, and sorted by abundance, prior chimera detection and OTU (operational taxonomic unit) determination at 97% sequence identity. Finally, original trimmed and high-quality sequences were mapped to OTUs at the 97% identity threshold to obtain one OTU table. The taxonomic affiliation of each OTU was obtained using the -*sintax* algorithm against the Ribosomal Database Project 16S rRNA training set version 18, with a confidence threshold of 80%. Rarity thresholds are typically defined as 0.1% or 0.01% of a taxon’s relative abundance (Pascoal et al., 2025). To determine the most appropriate cutoff for our study, we generated rank-abundance curves for the three soils (Fig. S1). These curves indicated that applying an OTU 0.1% threshold would overestimate the rare biosphere, particularly in the agricultural and forest soils. Therefore, we adopted an OTU relative abundance of 0.01% as the rarity threshold in this study.

The raw sequences associated with this study were deposited in the GenBank SRA database under BioProject accession number PNHJAC1367487.

### 2.5 Statistical analyses

One-way ANOVA (analysis of variance) was applied to determine whether the differences between treatments were significant (P < 0.05) using the *aov* function of the R package *stats* ver. 4.1.1 (R Core Team). When one-way ANOVA resulted in significant results, Tukey’s HSD (honest significance difference) post-hoc test was used for multiple comparisons of means at a 95 % confidence interval. The function *TukeyHSD* and the R packages *multcomp* ver. 1.4-23 (Hothorn et al., 2016) and *multcompView* ver. 0.1-9 (Graves et al.,2015) were used. Normality and heteroscedasticity of data were tested by the Kolmogorov–Smirnov (*ks*.*test* function, *stats* R package ver. 4.1.1) and Levene tests (*leveneTest* function, *car* R package ver. 3.1-1), respectively.

Venn diagrams were constructed with the R package *ggVennDiagram* ver. 1.5.4 (Duşa, 2024). To construct Venn diagrams, replicates from each treatment were merged with the function *merge*.*groups* of mothur ver. 1.48.3 (Schloss et al., 2009).

Data visualisations were conducted using the R package ggplot2 ver. 3.5.1 (Wickham, 2016) and CorelDRAW ver. 2020.

## 3 Results

### 3.1 Nutrient content of culture media

In general, macronutrient content in the culture media decreased in the following order: TSA > R2A > SEM> 1:20 R2A (Table S2). The TSA medium contained substantially higher nutrient levels than the other media. For example, it contained up to 50, 300, and 60 times more TOC, N, and P, respectively, than the other media. The nutrient content of each SEM mirrored that of the source soil (Table S1 and S2).

### 3.2 CFU counts

SEM yielded significantly higher bacterial CFU counts than the other tested media in the three soils. In contrast, TSA yielded the lowest CFU counts in agricultural and forest soils, while 1:20 R2A did so in contaminated soil (Fig. 1). CFU counts across media decreased with the type of soil in the following order: agricultural soil > forest soil > contaminated soil.

**Fig. 1.**
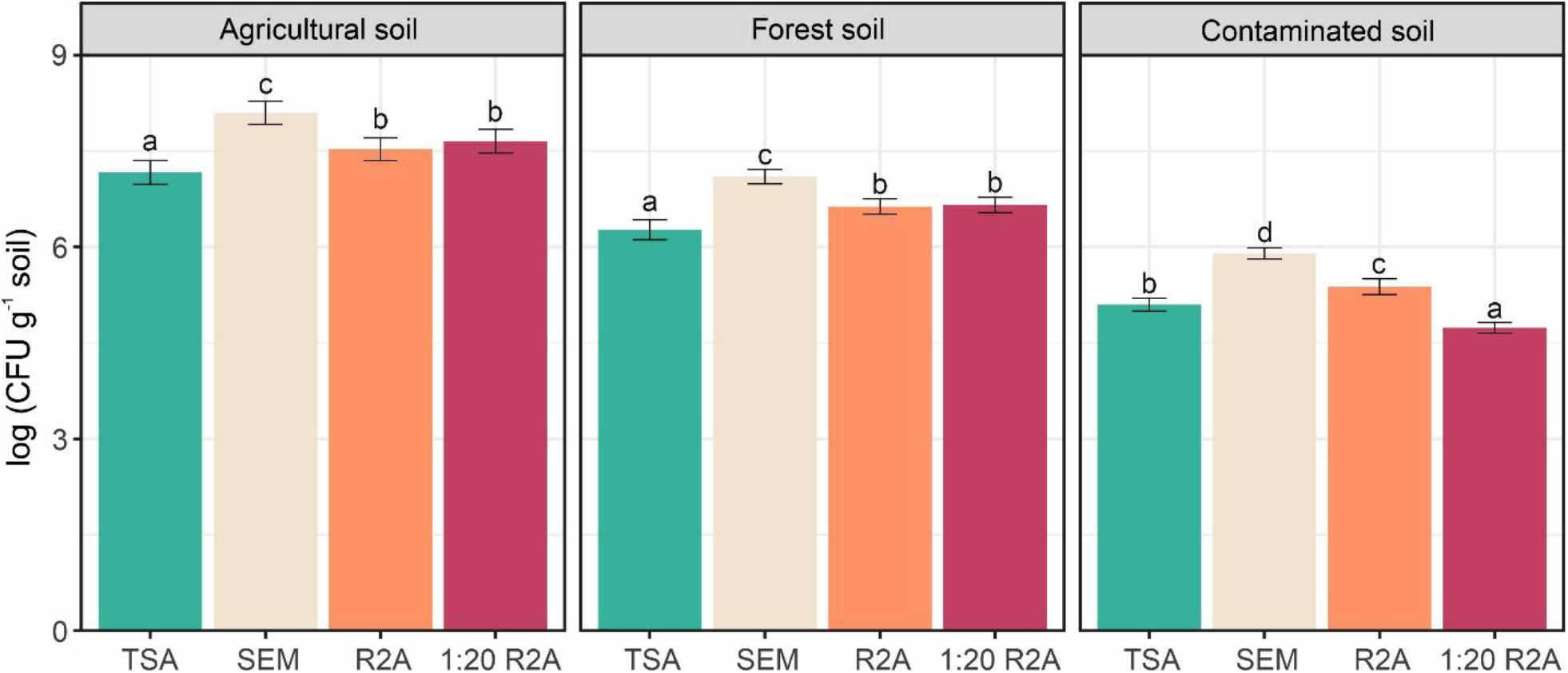
Colony-forming unit (CFU) counts for TSA (tryptic soy agar), SEM (soil extract medium), R2A (Reasoner’s 2A), and 1:20 R2A (Reasoner’s 2A diluted 20-fold) from agricultural, forest, and contaminated soils.

### 3.3 Culture-independent and culture-dependent richness and community taxonomic composition

In the three soils, culture-independent richness was much higher than that retrieved by culturing (Fig. 2). In general terms, culture dependent richness decreased with culture media in the following order: SEM > 1:20 R2A > R2A > TSA in the three soils. SEM yielded significantly greater richness in the three soils, whereas TSA recovered the lowest richness in the agricultural and forest soils. In the contaminated soil, no significant differences were observed between TSA, R2A, and 1:20 R2A (Fig 2).

**Fig. 2.**
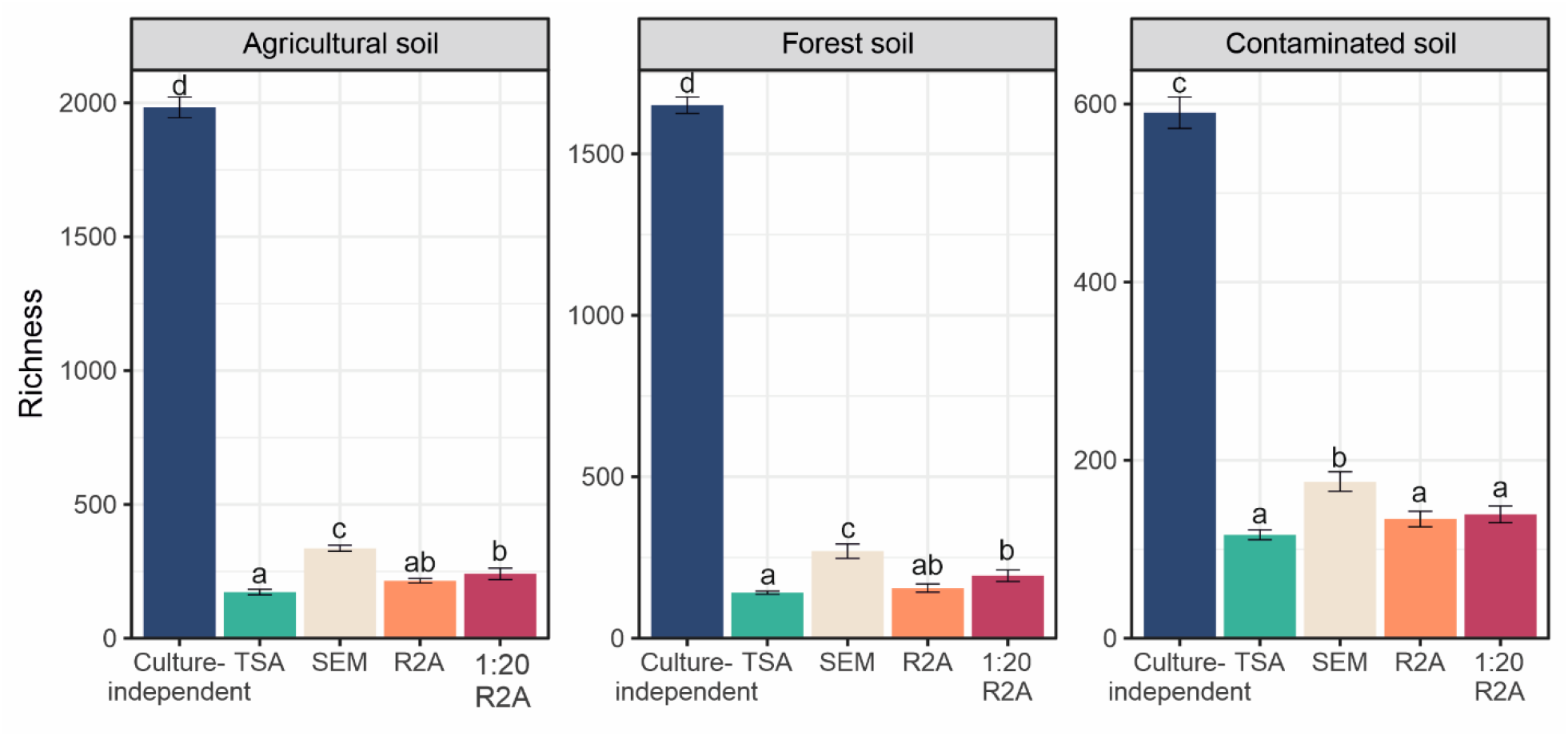
Culture-independent richness and culture-dependent richness using TSA (tryptic soy agar), SEM (soil extract medium), R2A (Reasoner’s 2A), and 1:20 R2A (Reasoner’s 2A diluted 20-fold) from agricultural, forest, and contaminated soils. For each soil, different letters above the standard deviation bars indicate significant differences (P < 0.05) according to ANOVA followed by Tukey’s HSD test.

The structure of the bacterial community was significantly affected by soil type and methodological approach (culture-independent vs. -dependent), with soil type being the most significant factor (Fig. S2). This was evidenced at taxonomic level. In the agricultural soil, culture-independent bacterial diversity was dominated by Chitinophagia, Actinobacteria, Betaproteobacteria, Cytophagia, and Bacilli at class level (Fig. 3). In this soil, culture-dependent diversity consisted at class level mainly of Gammaproteobacteria (*Stenotrophomonas* and *Pseudomonas* were the most abundant genera within this class), Bacilli (*Bacillus* and *Peribacillus*), and Actinobacteria (*Nocardioides*), with their relative proportions varying according to the culture medium (Fig. 3 and Table S3). SEM recovered the most diverse bacterial community (26 different classes from 11 phyla) and TSA the least diverse (19 classes from 8 phyla).

**Fig. 3.**
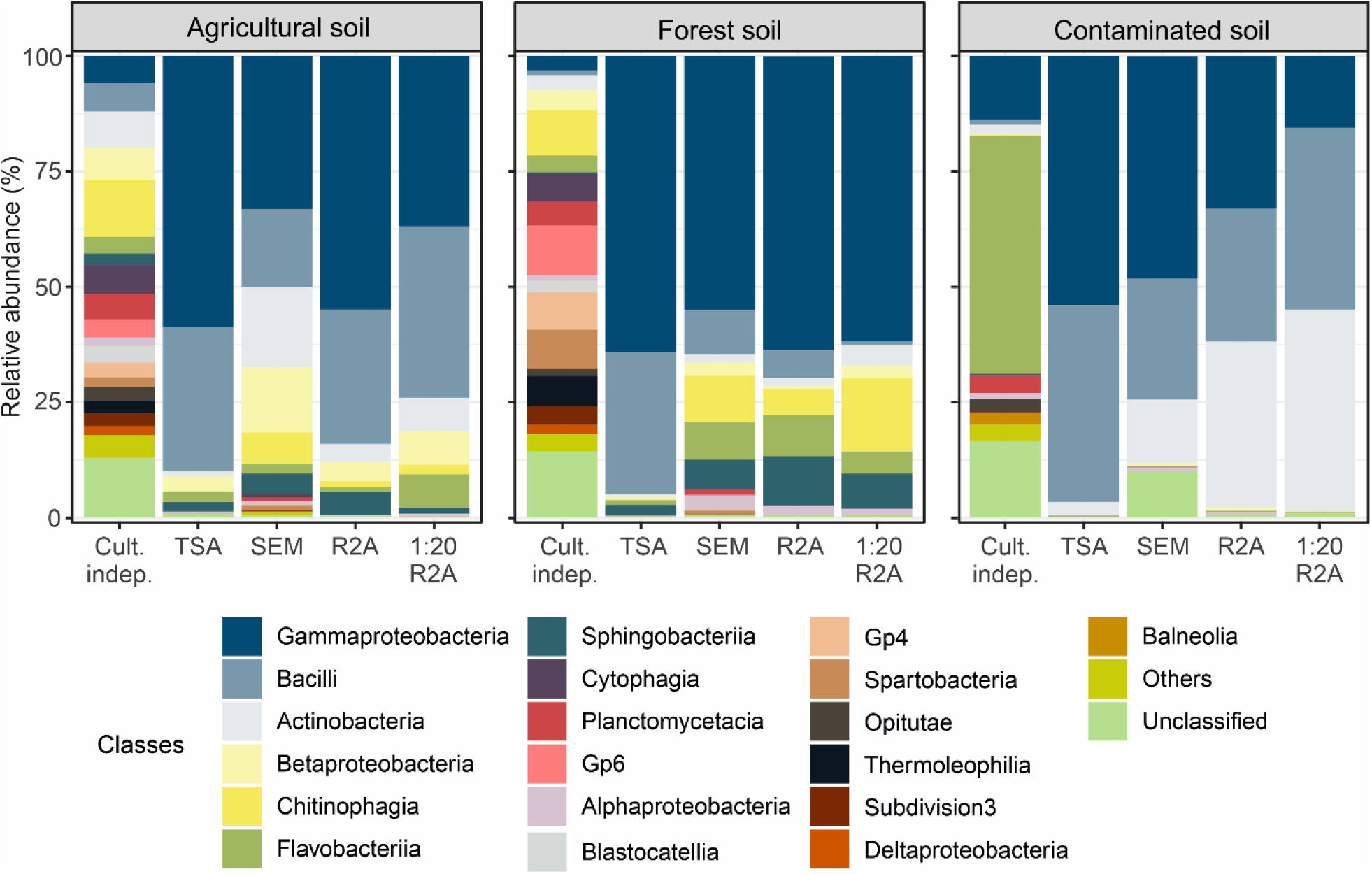
Taxonomic composition at class level of the culture-independent community and the culture-dependent community using TSA (tryptic soy agar), SEM (soil extract medium), R2A (Reasoner’s 2A), and 1:20 R2A (Reasoner’s 2A diluted 20-fold) from agricultural, forest, and contaminated soils.

Gp6, Chitinophagia, Spartobacteria, Gp4, Thermoleophilia, Cytophagia, and Planctomycetacia were the classes dominating culture-independent diversity in the forest soil (Fig. 3). The culture-dependent diversity captured with the different media was dominated by Gammaproteobacteria (*Pseudomonas* and *Serratia*), Bacilli (*Peribacillus*), and Chitinophagia (*Chitinophaga*) (Fig. 3 and Table S4). SEM recovered the most taxonomically diverse community (26 different classes from 12 phyla), while R2A recovered the least diverse (17 different classes from 10 phyla).

Taxonomic composition of the culture-independent community in the contaminated soil was clearly dominated by the class Flavobacteriia, with other major classes being Gammaproteobacteria and Planctomycetacia (Fig. 3). Culture-dependent diversity was dominated by Gammaproteobacteria (*Pseudomonas* and *Lysobacter*), Bacilli (*Mesobacillus* and *Bacillus*), and Actinobacteria (*Streptomyces* and *Nocardioides*) (Fig. 3 and Table S5). Consistent with the results from the other soils, SEM retrieved the most diverse bacterial community (22 different classes from 12 phyla), while TSA showed the lowest (17 different classes from 10 phyla).

### 3.4 Richness recovered by culture-dependent approaches relative to culture-independent community

We quantified the proportion of OTUs recovered by culturing relative to those detected by the culture-independent approach (Fig. 4 and Table 1 and S6). SEM yielded significantly higher proportions of OTU recovering than the other media. In this way, this medium was able to recover up to 13%, 9%, and 18% of the culture-independent OTUs in agricultural, forest, and contaminated soils, respectively (Table 1). Further, in the agricultural soil, R2A and 1:20 R2A recovered a higher proportion of the culture-independent diversity compared to TSA. However, this pattern was not observed in the forest and contaminated soils. When the culturable diversity of the four soil replicates was considered altogether, rather than separately, and compared with the combined culture-independent diversity of the same replicates, SEM proved to be the most effective method for recovering the culture-independent community, with recovery rates of 21%, 16%, and 26% for agricultural, forest, and contaminated soils, respectively (Table 1). This also demonstrates the value of replication in microbial isolation within culture-dependent studies.

**Table 1.** Percentages of OTUs recovered through the culture-dependent approach using TSA (tryptic soy agar), SEM (soil extract medium), R2A (Reasoner’s 2A), and 1:20 R2A (Reasoner’s 2A diluted 20-fold) in agricultural, forest, and contaminated soils, relative to those obtained through the culture-independent approach. Data are presented considering replicates both separately and jointly. For the part of the table showing replicates independently, values represent mean ± standard deviation (n = 4) and, for each medium, different letters indicate significant differences (P < 0.05) according to ANOVA followed by Tukey’s HSD test.

| Culture medium | Agricultural soil | Forest soil | Contaminated soil |
| --- | --- | --- | --- |
| Considering replicates independently |  |  |  |
| TSA (%) | 6.09 ± 0.22 a | 5.27 ± 0.41 a | 11.07 ± 1.57 a |
| SEM (%) | 12.66 ± 0.41c | 8.84 ± 1.70 b | 18.14 ± 1.18 b |
| R2A (%) | 8.09 ± 0.15 b | 3.64 ± 0.79 a | 12.51 ± 1.00 a |
| 1:20 R2A (%) | 9.05 ± 1.05 b | 4.89 ± 0.29 a | 12.67 ± 1.31 a |
| Considering replicates jointly |  |  |  |
| TSA (%) | 11.23 | 10.96 | 15.54 |
| SEM (%) | 21.04 | 15.92 | 25.89 |
| R2A (%) | 14.40 | 6.13 | 18.74 |
| 1:20 R2A (%) | 16.49 | 8.23 | 19.48 |
| All media (%) | 30.74 | 22.23 | 35.38 |

**Fig. 4.**
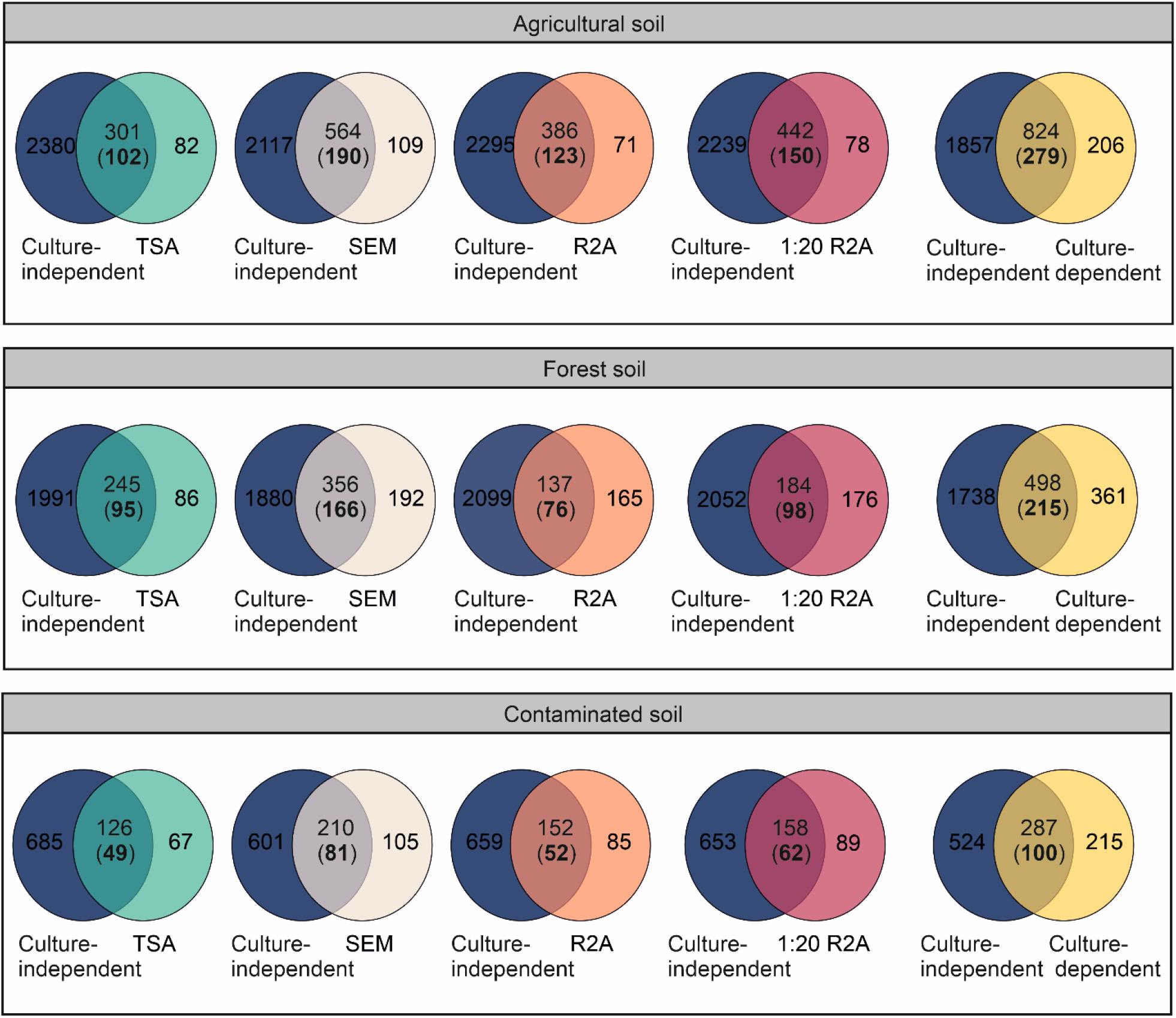
Venn diagram showing shared and exclusive OTUs for the culture-independent and culture-dependent fractions, as well as for each culture medium. Numbers in black in parentheses within the shared fraction represent rare OTUs, i.e., those with a relative abundance below 0.01%.TSA, tryptic soy agar; SEM, soil extract medium; R2A, Reasoner’s 2A;1:20 R2A, Reasoner’s 2A diluted 20-fold.

### 3.5 Culture-independent and culture-dependent rare taxonomic community

In the agricultural soil, culture-independent rare community consisted of 1,163 OTUs across the four replicates. Compared with the taxonomic composition of the general culture-independent community, the rare community was richer in Chitinophagia, Betaproteobacteria, and Blastocatellia, and poorer in Planctomycetacia and Actinobacteria (Fig. S3 and Fig. 3). The culture-independent rare community in the forest soil comprised 1,320 OTUs. Respect to the taxonomic composition of the general community, the rare community was richer in Gp4, Gp6, Spartobacteria, Thermoleophilia, and Chitinophagia, and poorer in Actinobacteria, Planctomycetacia, Sphingobacteriia, and Verrucomicrobiae (Fig. S4). In the contaminated soil, the culture-independent rare community comprised 617 OTUs, with its taxonomic composition being richer in Flavobacteriia and Opitutae, and poorer in Planctomycetacia, Bacilli, Actinobacteria, and Alphaproteobacteria in comparison with the general community (Fig. S4).

We then analyzed the number of OTUs retrieved by culturing that were members of the rare culture-independent community. Across the three soils, SEM isolated a significantly greater number of rare culture-independent OTUs, with the differences being most pronounced compared to TSA (Fig 4 and Table 2 and S6). However, when the proportions of rare OTUs were calculated relative to the total number of isolated OTUs, no significant differences were detected between culture media (Table 2). In general, an important fraction of the OTUs retrieved by culturing were members of the rare community. In the agricultural and contaminated soils, between 30 and 40% of the retrieved OTUs were classified as rare, while this proportion increased up to ~56% in the forest soil when using the R2A medium (Table 2).

**Table 2.**
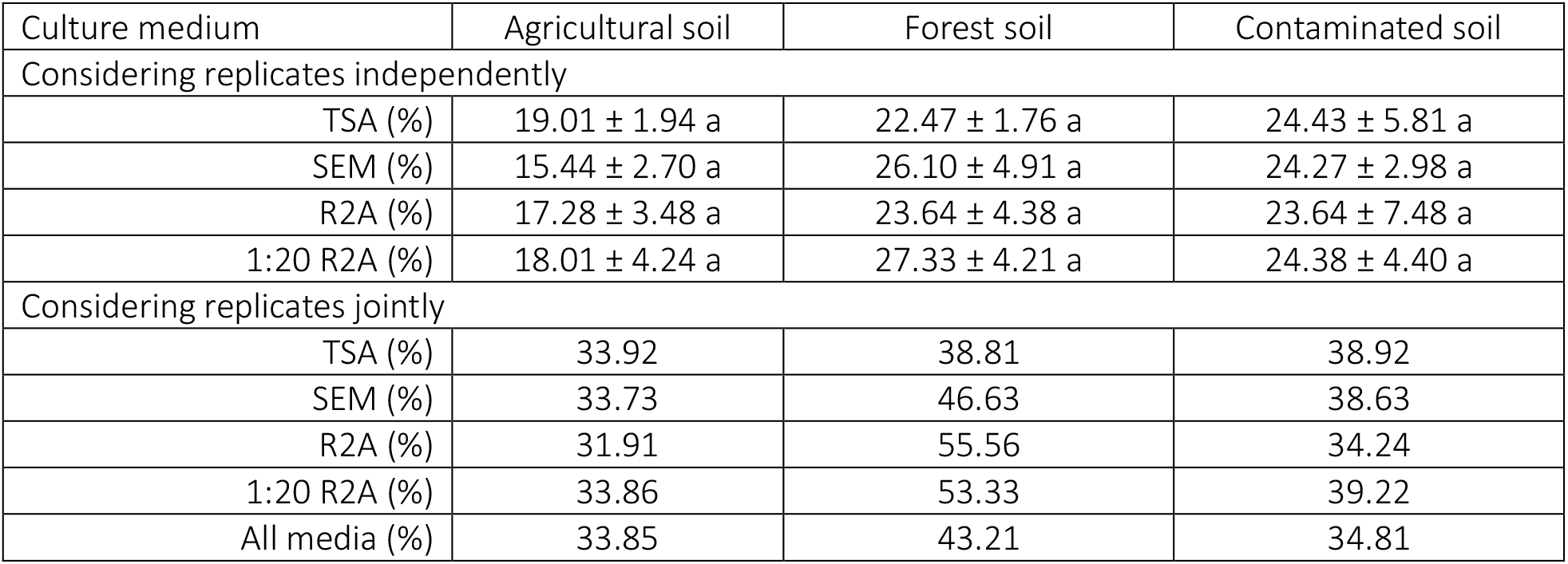
Percentages of rare OTUs relative to total OTUs recovered through the culture-dependent approach using TSA (tryptic soy agar), SEM (soil extract medium), R2A (Reasoner’s 2A), and 1:20 R2A (Reasoner’s 2A diluted 20-fold) in agricultural, forest, and contaminated soils. Data are presented considering replicates both separately and jointly. For the part of the table showing replicates independently, values represent mean ± standard deviation (n = 4) and, for each medium, different letters indicate significant differences (P < 0.05) according to ANOVA followed by Tukey’s HSD test.

| Culture medium | Agricultural soil | Forest soil | Contaminated soil |
| --- | --- | --- | --- |
| Considering replicates independently |  |  |  |
| TSA (%) | 19.01 ± 1.94 a | 22.47 ± 1.76 a | 24.43 ± 5.81 a |
| SEM (%) | 15.44 ± 2.70 a | 26.10 ± 4.91 a | 24.27 ± 2.98 a |
| R2A (%) | 17.28 ± 3.48 a | 23.64 ± 4.38 a | 23.64 ± 7.48 a |
| 1:20 R2A (%) | 18.01 ± 4.24 a | 27.33 ± 4.21 a | 24.38 ± 4.40 a |
| Considering replicates jointly |  |  |  |
| TSA (%) | 33.92 | 38.81 | 38.92 |
| SEM (%) | 33.73 | 46.63 | 38.63 |
| R2A (%) | 31.91 | 55.56 | 34.24 |
| 1:20 R2A (%) | 33.86 | 53.33 | 39.22 |
| All media (%) | 33.85 | 43.21 | 34.81 |

In the agricultural soil, the taxonomic composition of the isolated rare community was comprised of 20 different classes, being the most abundant Bacilli, Actinobacteria and Gammaproteobacteria (Fig. 5). SEM retrieved the most diverse community, with 58 different genera (Table S7). At genus level, *Bacillus* predominated in TSA, R2A, and 1:20 R2A among rare diversity, while *Nocardioides* did so in SEM. Other abundant genera comprising rare community were *Solibacillus, Paenibacillus*, and *Lysinibacillus* in TSA; *Chitinophaga, Xanthomonas* and *Pseudoxanthomonas* in SEM; *Pedobacter, Microbacterium*, and *Agromyces* in R2A; and *Nocardioides, Xanthomonas*, and *Williamsia* in 1:20 R2A (Table S7).

**Fig. 5.**
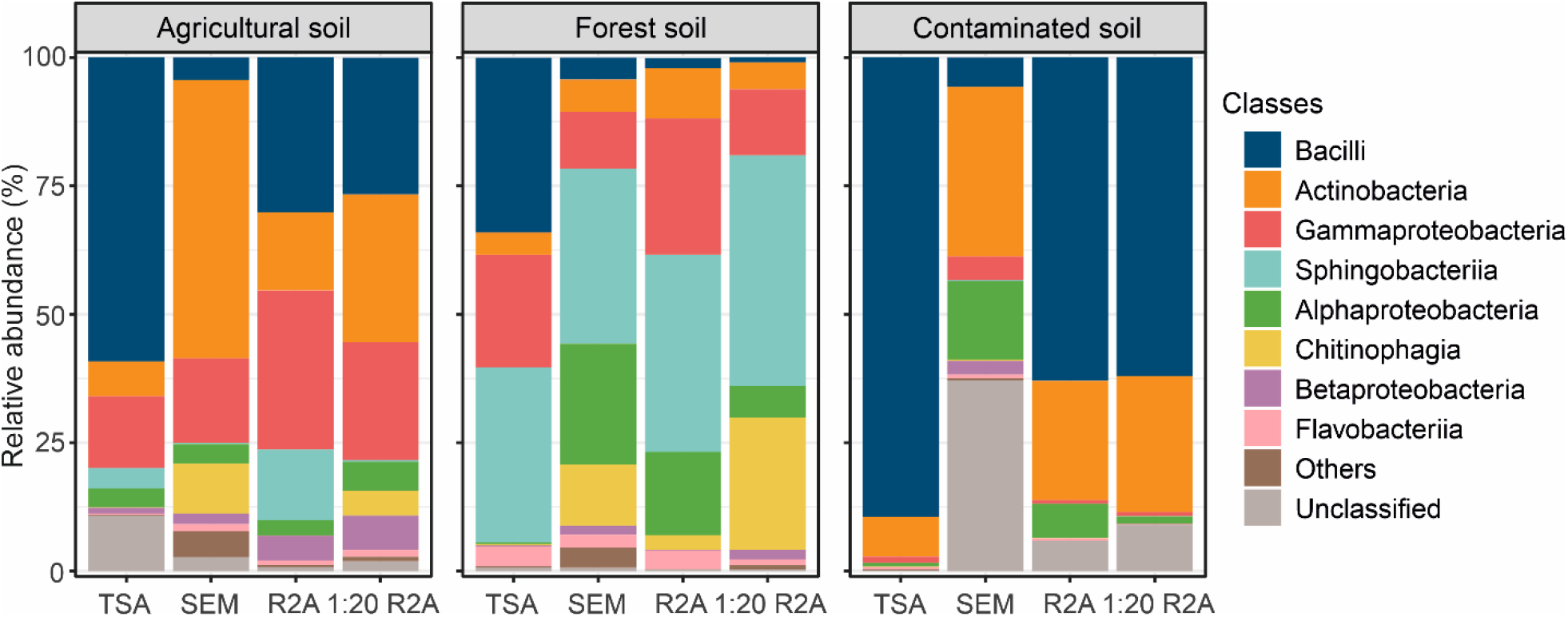
Taxonomic composition at class level of the rare bacterial fraction isolated using TSA (tryptic soy agar), SEM (soil extract medium), R2A (Reasoner’s 2A), and 1:20 R2A (Reasoner’s 2A diluted 20-fold) from agricultural, forest, and contaminated soils.

In the forest soil, the rare community recovered across the four media comprised 19 classes, with Sphingobacteriia, Gammaproteobacteria, Alphaproteobacteria, Bacilli, and Chitinophagia dominating (Fig. 5). SEM retrieved the most diverse community, with 53 different genera (Table S8). Within the rare community obtained from the TSA medium, the genera *Sphingobacterium* and *Sporosarcina* were predominant. Rare communities isolated using SEM and R2A were largely composed of *Pedobacter* and *Phyllobacterium*, while the rare community obtained with 1:20 R2A was dominated by *Pedobacter, Chitinophaga*, and *Pseudomonas* (Table S8).

In the contaminated soil, the rare community recovered across the four media encompassed 15 different classes, with Bacilli and Actinobacteria being the most abundant (Fig. 5). SEM and 1:20 R2A recovered a more diverse rare community than the other media, with 28 different genera (Table S9). In TSA, rare community was dominated at genus level by *Bacillus* and *Solibacillus*; in R2A, by *Bacillus, Fictibacillus*, and *Streptomyces*; and in 1:20 R2A, by *Bacillus, Neobacillus*, and *Streptomyces*. In contrast, the rare diversity retrieved by SEM was dominated by *Janibacter* and *Georgenia*. It is worth noting that 67% of the rare taxa retrieved by SEM could not be classified at genus level (Table S9).

### 3.6 Richness and taxonomic composition of the bacterial fraction captured only by culturing

A significant number of OTUs retrieved by the culture-dependent approach were not present in the culture-independent dataset (Fig. 4 and Table S6). SEM retrieved a significantly higher number of unique OTUs than the other media in the three soils, and R2A and 1:20 R2A isolated more unique OTUs than TSA in the forest and contaminated soils, but not in the agricultural soil (Table S6). When the four replicates per soil are collectively considered, SEM recovered 109, 192, and 105 unique culturable OTUs, representing approximately 7%, 9%, and 12% of the OTUs detected by the culture-independent approach in the agricultural, forest, and contaminated soils, respectively.

In the agricultural soil, the taxonomic composition of the unique culturable fraction retrieved by the four media was comprised 21 different classes, with Bacilli, Alphaproteobacteria, Actinobacteria, and Sphingobacteriia being the most abundant (Fig. 6). SEM was the medium retrieving the most diverse unique culturable fraction with 32 different genera, while R2A, TSA, and 1:20 R2A retrieved 23, 21 and 19 different genera, respectively (Table S10). The unique culturable fraction retrieved by TSA and R2A was clearly biased towards *Bacillus*. However, the diversity captured by 1:20 R2A and SEM was more evenly distributed among the most abundant genera. *Staphylococcus, Microbacterium*, and *Bacillus* predominated among the unique culturable diversity recovered by 1:20 R2A, whereas *Olivibacter, Fictibacillus*, and *Nocardioides* were dominant among the culturable diversity obtained via SEM. In this medium, ~47% of the unique culturable sequences could not be taxonomically affiliated at genus level (Table S11).

**Fig. 6.**
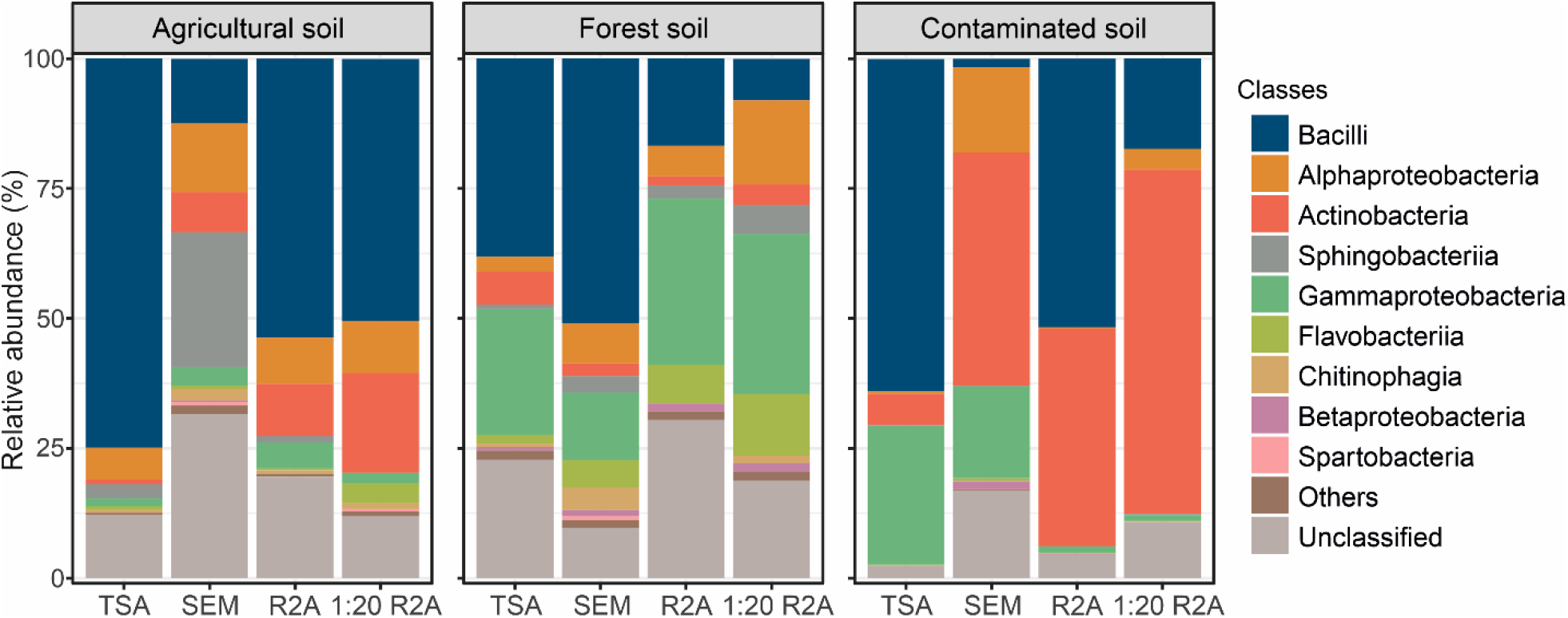
Taxonomic composition at class level of the unique bacterial fraction isolated using TSA (tryptic soy agar), SEM (soil extract medium), R2A (Reasoner’s 2A), and 1:20 R2A (Reasoner’s 2A diluted 20-fold) from agricultural, forest, and contaminated soils.

In the forest soil, the unique culturable fraction retrieved with the four culture media was composed of 24 different classes (Fig. 6). Bacilli, Gammaproteobacteria, Alphaproteobacteria, and Flavobacteriia were the most abundant. The number of unique culturable genera decreased with the culture medium in the following order: SEM (65) > 1:20 R2A (58) > R2A (48) > TSA (34) (Table 11). Within the unique culturable diversity retrieved by each culture medium, *Bacillus, Pseudomonas*, and *Sporosarcina* predominated on TSA; *Bacillus, Pseudomonas*, and *Flavobacterium* on SEM; *Pseudomonas, Lysobacter*, and *Paenibacillus* on R2A; and *Pseudomonas, Lysobacter*, and *Flavobacterium* on 1:20 R2A (Table S11).

In the contaminated soil, the unique culturable fraction recovered across the four culture media comprised 10 classes, with the most abundant being Actinobacteria, Bacilli, Gammaproteobacteria, and Alphaproteobacteria (Fig. 6). The number of different unique culturable genera decreased with the culture medium in the following order: 1:20 R2A (40) > SEM (30) > R2A (30) > TSA (25) (Table S12). *Bacillus, Lysobacter*, and *Solibacillus* were the most abundant genera conforming unique culturable fraction in TSA; *Dietzia, Pseudactinotalea*, and *Pseudomonas* in SEM; *Paenibacillus, Bacillus*, and *Streptomyces* in R2A; and *Streptomyces, Cellulosimicrobium*, and *Pseudactinotalea* in 1:20 R2A (Table S12).

## 4 Discussion

Our culture-dependent approach revealed that 31%, 22%, and 35% of the OTUs in the culture-independent dataset were recoverable through culturing in agricultural, forest, and contaminated soils, respectively. These percentages are much higher than those reported in other studies assessing culture-dependent communities via culture plating and subsequent colony picking (Pascual et al., 2016; Siles et al., 2022; Stefani et al., 2015; Youseif et al., 2021). Although our culture-dependent approach was more intensive than those used in previous studies, the observed differences may also result from the loss of diversity that occurs when colony picking is employed. Our recovery percentages were also higher than those reported by Shade et al. (2012) using the same experimental approach, which can be attributed to our more intensive culture-dependent effort. Recovery success was higher in agricultural and contaminated soils than in forest soil, which is seen in relation to the higher degree of human disturbance in the first two soils. Disturbed soils likely favor fast-growing, copiotrophic bacteria and pollutant-degrading taxa that readily form colonies, whereas forest soils may support communities dominated by slow-growing oligotrophs and specialists that are more difficult to cultivate (Carbonetto et al., 2014; Dragone et al., 2024). Additionally, when considering individual soil replicates, the mean proportion of culturable OTUs was lower than that obtained by combining all four replicates, nearly doubling the recovery rate when pooled. This highlights the importance of including replicates in bacterial isolation studies, given the high spatial heterogeneity of soil bacterial communities (Grzyb et al., 2022).

Among the four media tested, SEM proved to be the most effective for recovering a more abundant (CFU counts), richer (OTU number, richness), and more taxonomically diverse community across the three soils. Accordingly, SEM recovered 21%, 16%, and 26% of the OTUs detected by metataxonomics in the agricultural, forest, and contaminated soils, respectively. This finding supports our initial hypothesis, as we anticipated that a nutrient-poor medium with a chemical composition relatively similar to that of soil would be the most effective for recovering a richer and more taxonomically diverse community. Our initial assumption was based on previous studies successfully isolating new bacteria using SEM (Mandakovic et al., 2018; Nguyen et al., 2018) and on evidence that oligotrophic or diluted media can facilitate recovery of a more diverse microbial community (Dai et al., 2025; Davis et al., 2005; Wang et al., 2020). Nonetheless, the physiological mechanisms causing the enhanced growth of diverse microbes in low-nutrient media, and the inhibition of certain taxa under high-nutrient conditions, remain poorly understood. One possible explanation for these concentration-dependent effects is that growth media at high concentrations may contain substantial amounts of inhibitory compounds, which are reduced to non-inhibitory levels in more dilute formulations (Bartelme et al., 2020). Additionally, our findings suggest that nutrient composition may be more important than nutrient concentration for isolating more diverse communities of both abundant and rare soil bacteria (Bartelme et al., 2020). TSA is a rich medium whose C sources are mainly derived from digests of casein and soybean, providing a high concentration of peptides, amino acids, and some simple carbohydrates. R2A contains glucose and soluble starch as direct carbon sources, supplemented by yeast extract, proteose peptone, and casamino acids that supply additional amino acids and small peptides. SEM is rich in complex organic compounds such as humic and fulvic acids, aromatic compounds, polysaccharides, and a variety of organic acids, along with trace elements. In this way, while TSA and R2A provide readily metabolizable C and N sources for general or oligotrophic bacteria, SEM offers a chemically diverse, natural substrate spectrum that more accurately reflects the microbial nutrient environment in soil ecosystems, supporting the growth of a higher bacterial community (Nguyen et al., 2018).

SEM not only sustained the highest total culture-dependent richness but also retrieved the largest absolute number of rare OTUs in the three soils. Despite differences in absolute numbers, the relative proportion of rare OTUs (i.e., relative to the total number of isolated OTUs) remained relatively stable across media, averaging ~34% in agricultural and contaminated soils and ~43% in forest soil. The higher relative proportion of rare bacteria recovered from forest soil reflects the larger size of the culture-independent rare community in this soil compared to the others. These findings suggest that while medium composition and nutrient content influence the absolute recovery of rare taxa, the relative representation of rare OTUs within the cultured community is largely maintained, reflecting intrinsic patterns of culturability and competitive dynamics among soil bacteria (Shade et al., 2012). SEM’s effectiveness likely arises from its moderately poor but chemically diverse nutrient environment, which supports both fast-growing (copiotrophic) and slow-growing (oligotrophic) taxa. In contrast, very rich media such as TSA preferentially favor fast-growing taxa, potentially suppressing rare species, whereas extremely poor media (e.g., 1:20 R2A) may limit overall growth, preventing some rare taxa from forming colonies. Our elemental analysis of the culture media supported this argument, showing that nutrient content decreased as follows: TSA >> R2A > SEM > 1:20 R2A. Therefore, SEM achieves a balance: it supports sufficient growth to allow a high total OTU richness, which naturally increases the absolute number of rare OTUs. The effectiveness of SEM to recover more OTUs is thus proportional to both abundant and rare taxa. The fact that relative proportions remain similar across media also suggest that rare taxa maintain their ecological status (as rare) regardless of nutrient conditions, implying that rarity is a property of the community that is largely independent of the medium used for culturing.

The rare culture-dependent diversity across the four media was primarily composed of Proteobacteria, Actinobacteria, Bacteroidetes, and Firmicutes across the three soils. A detailed analysis of the genera recovered from each medium revealed a consistent pattern, which we linked to nutrient availability and microbial ecological strategies. In this way, Firmicutes spore-forming genera (e.g., *Bacillus, Paenibacillus, Solibacillus, Lysinibacillus, Fictibacillus, Neobacillus*) were consistently favored by TSA and R2A. Although R2A is generally regarded as an oligotrophic medium, our analysis showed that it contains a much higher nutrient content than SEM and 1:20 R2A. Additionally, we noticed that the richer the soil was in spore-forming genera, as was the case with agricultural and contaminated soils, the more such taxa were isolated. Their dominance likely reflects the ability of spores to withstand environmental stress in soil and rapidly germinate under favorable nutrient conditions, providing a competitive advantage over slower-growing taxa. In contrast, SEM, with its chemically diverse but moderately poor nutrient composition resembling soil, supported a rare community dominated by non-spore-forming genera such as *Nocardioides, Chitinophaga, Xanthomonas, Janibacter*, and *Georgenia*, likely because it reduces competitive exclusion by fast-growing spore-formers. Both spore-forming and non-spore-forming taxa are components of the soil rare biosphere; however, SEM preferentially isolated non-spore-forming taxa, which is particularly relevant for understanding soil functioning, as these taxa are more likely to be metabolically active in situ (Chen et al., 2020; Liang et al., 2019). In contrast, spore-forming taxa often exist as dormant spores and may not contribute directly to ongoing soil processes under natural conditions. Surprisingly, under extreme nutrient limitation, as in 1:20 R2A, the relative abundance of spore-forming genera such as *Bacillus* and *Streptomyces* again increased. This likely reflects their ability to germinate from dormant spores when trace nutrients are available and to utilize scarce or recalcitrant compounds more efficiently than non-spore-formers, maintaining a competitive advantage under oligotrophic conditions (Stewart and Setlow, 2013; Xu and Vetsigian, 2017). Together, these observations indicate that spore-forming taxa appear to be preferentially recovered under both nutrient-rich and nutrient-poor extremes, whereas soil-like media such as SEM provide conditions that facilitate the cultivation of a broader spectrum of rare, non-spore-forming taxa.

SEM not only reduced bias toward spore-forming taxa but also facilitated the isolation of rare taxa from underrepresented phyla in culture-dependent studies (Lewis et al., 2020), a result that was observed to some extent for 1:20 R2A as well. For example, SEM was able to recover members of the rare biosphere belonging to Acidobacteria (*Edaphobacter*, retrieved from agricultural soil; *Terriglobus*, forest soil), Planctomycetes (*Singulisphaera*, agricultural and forest soils, *Gemmata*, agricultural and forest soils; *Schlesneria*; agricultural soil; *Blastopirellula*, contaminated soil), or Verrucomicrobia (*Verrucomicrobium*, agricultural soil; Spartobacteria_genera_incertae_sedis, agricultural, forest and contaminated soils) phyla. These rare taxa are known to occupy specialized niches, metabolize recalcitrant compounds, and produce secondary metabolites that influence microbial communities and plant–microbe interactions, among other functions (Ivanova et al., 2016; Kielak et al., 2016; Nixon et al., 2019). Moreover, they are considered a functional “seed bank,” harboring genetic and metabolic diversity that can be activated under changing environmental conditions, thereby sustaining key ecosystem functions and enhancing resilience (Lennon and Jones, 2011; Pascoal et al., 2021). Identifying SEM as an effective approach for isolating these rare taxa represents a promising opportunity to advance our understanding of their functional and ecological roles.

A considerable number of OTUs detected through the culture-dependent approach were not found in the culture-independent dataset. Although this discrepancy could partly result from biases in DNA extraction or PCR amplification, it is more likely that culturing retrieved bacterial taxa that are below the detection limit of the culture-independent approach, even though rarefaction curves indicated that our sequencing depth was sufficient to capture the overall soil bacterial diversity (Fig. S4). This phenomenon has been reported previously, with the number of unique culturable taxa varying according to cultivation effort (Hinsu et al., 2021; Lee et al., 2016; Siles et al., 2022). In our study, the culture-dependent approach revealed 206, 361, and 215 unique OTUs in the agricultural, forest, and contaminated soils, respectively. When comparing these unique OTUs with culture-independent richness from each soil, we found that up to ~8%, 16%, and 27% of the bacterial community in the agricultural, forest, and contaminated soils, respectively, could only be detected through culturing. Our proportions were in line with those obtained by Shade et al. (2012), which were of ~14%. In our study, if unique culturable OTUs, potentially rare taxa, are combined with those culturable OTUs identified as rare in the culture-independent dataset, then up to 485, 576, and 315 OTUs of the rare biosphere in agricultural, forest, and contaminated soils, respectively, were recovered by our culture-dependent approach. This confirms that culturing is more effective than initially expected in capturing members of the rare biosphere, serving not only to provide isolates for further studies but also as a highly reliable tool to complement metataxonomic data for a more comprehensive description of soil bacterial diversity (Shade et al., 2012).

A key strength of our study compared to previous works is that we evaluated how the type of culture media influences the recovery of unique culturable bacteria. SEM was the medium that most effectively isolated higher richness in the three soils. Overall, when considering the four media, increased nutrient concentration in the media corresponded to a decrease in richness of this fraction. Further, SEM was the medium that recovered the highest taxonomic diversity (number of different genera) and also sustained the highest relative abundance of unclassified sequences (>40%) at the genus level in agricultural and contaminated soils. While high proportions of unclassified sequences can result from (i) short sequence lengths or (ii) incomplete reference databases (Rachid et al., 2013), in this study most sequences were successfully classified at higher taxonomic levels, including kingdom (Bacteria) and phylum. Therefore, the high proportion of sequences lacking genus-level classification indicates that SEM supports the growth of novel or poorly described taxa that remain unresolved at fine taxonomic resolution (Delgado-Baquerizo, 2019), suggesting that low-nutrient, soil-derived media favor the recovery of phylogenetically diverse, yet underexplored, members of the rare biosphere.

Contrary to our initial hypothesis, the unique culturable fraction was not taxonomically exclusive. We observed a mixed pattern: while several rare genera were found exclusively within the unique culturable fraction, others overlapped with taxa identified as rare and present in the culture-independent dataset. For instance, in the agricultural forest, unique culturable taxa were affiliated with the genus *Bacillus*, which was also represented among rare taxa in the culture-independent dataset. A similar pattern was observed for the genus *Pedobacter* in the forest soil. This is explained by the fact that each genus comprises multiple taxa (OTUs) with varying levels of rarity (Bickel and Or, 2021). Alternatively, we found a number of genera that were exclusive of the unique culturable fraction in each soil (Tables S10-S12). In both agricultural and forest soils, SEM was the most effective method for retrieving taxa affiliated with genera absent from the culture-independent dataset. For instance, SEM in the agricultural soil retrieved unique taxa belonging to genera such as *Devosia, Aeromicrobium, Nemorincola, Edaphobacter, Yinghuangia*, and *Dyella*, among others. In the forest soil, unique genera included *Sphingobium, Steroidobacter, Comamonas, Microbacterium, Lysinibacillus, Methylotenera, Rhodanobacter*, and *Streptosporangium*, among many others. Our analyses demonstrate that these taxa are rare members of the soil microbial community. Many of these genera are known for important functional traits in soil ecosystems, including the degradation of complex organic compounds (e.g., *Sphingobium, Steroidobacter, Comamonas*), nutrient cycling (e.g., *Microbacterium, Rhodanobacter*), plant growth promotion (e.g., *Lysinibacillus*), and tolerance to environmental stressors (Feller et al., 2021; Green et al., 2012; Mitra et al., 2020; Zhang et al., 2024). This highlights the potential ecological significance of the unique culturable fraction, which can capture functionally relevant taxa that are often overlooked by culture-independent approaches.

## 5 Conclusions

Intensive culturing can recover a substantial fraction (at least 30%) of soil bacterial diversity, including members of the rare biosphere, and this recovery markedly increases with repeated soil isolations. Among the media tested, SEM, characterized by its low nutrient content and chemical composition resembling that of soil, proved to be the most effective, yielding higher CFU counts, greater richness, and broader taxonomic diversity across the three studied soils. Although the relative proportion of rare OTUs among total isolates did not differ substantially between culture media, SEM provided enhanced access to the rare biosphere, as evidenced by the increased richness and taxonomic diversity of rare taxa recovered, representing a community not biased toward spore-forming bacteria. Culturing also captured a significant fraction of soil bacterial diversity that remained below the detection threshold of 16S rRNA metabarcoding, particularly when using SEM. This unique culturable fraction, potentially part of the soil rare biosphere, was taxonomically diverse, included a high proportion of unclassified sequences, and comprised both exclusive and overlapping taxa relative to the culture-independent community. Overall, these findings highlight the value of low-nutrient, soil-like media for cultivating taxonomically diverse bacteria of the rare soil biosphere, thereby complementing metabarcoding approaches and opening new opportunities to explore their ecological roles and contributions to soil functioning.

## Credit Author Statement

José A. Siles: Conceptualization, Methodology, Formal analysis, Investigation, Writing - Original Draft, Visualization. Norman Terry: Resources, Funding acquisition.

## Acknowledgements

This work was supported by the UC Berkeley Grant number 51719.

## References

Aanderud, Z.T., Jones, S.E., Fierer, N., Lennon, J.T., 2015. Resuscitation of the rare biosphere contributes to pulses of ecosystem activity. Front Microbiol 6, 24. 10.3389/fmicb.2015.00024.

Anthony, M.A., Bender, S.F., van der Heijden, M.G.A., 2023. Enumerating soil biodiversity. Proceedings of the National Academy of Sciences 120, e2304663120. doi:10.1073/pnas.2304663120.

Bartelme, R.P., Custer, J.M., Dupont, C.L., Espinoza, J.L., Torralba, M., Khalili, B., Carini, P., 2020. Influence of Substrate Concentration on the Culturability of Heterotrophic Soil Microbes Isolated by High-Throughput Dilution-to-Extinction Cultivation. mSphere 5, e00024–00020. 10.1128/mSphere.00024-20.

Bickel, S., Or, D., 2021. The chosen few—variations in common and rare soil bacteria across biomes. ISME J 15, 3315–3325. 10.1038/s41396-021-00981-3.

Carbonetto, B., Rascovan, N., Álvarez, R., Mentaberry, A., Vázquez, M.P., 2014. Structure, Composition and Metagenomic Profile of Soil Microbiomes Associated to Agricultural Land Use and Tillage Systems in Argentine Pampas. PLoS One 9, e99949. 10.1371/journal.pone.0099949.

Carini, P., 2019. A “Cultural” Renaissance: Genomics Breathes New Life into an Old Craft. mSystems 4, e00092–00019. 10.1128/mSystems.00092-19.

Chen, Q.-L., Ding, J., Zhu, D., Hu, H.-W., Delgado-Baquerizo, M., Ma, Y.-B., He, J.-Z., Zhu, Y.-G., 2020. Rare microbial taxa as the major drivers of ecosystem multifunctionality in long-term fertilized soils. Soil Biology and Biochemistry 141, 107686. 10.1016/j.soilbio.2019.107686.

Dai, J., Ouyang, Y., Gupte, R., Liu Xiao Jun, A., Li, Y., Yang, F., Chen, S., Provin, T., Van Schaik, E., Samuel James, E., Jayaraman, A., Zhou, A., de Figueiredo, P., Zhou, J., Han, A., 2025. Microfluidic droplets with amended culture media cultivate a greater diversity of soil microorganisms. Appl Environ Microbiol 91, e01794–01724. 10.1128/aem.01794-24.

Davis, K.E.R., Joseph, S.J., Janssen, P.H., 2005. Effects of Growth Medium, Inoculum Size, and Incubation Time on Culturability and Isolation of Soil Bacteria. Appl Environ Microbiol 71, 826–834. 10.1128/aem.71.2.826-834.2005.

Delgado-Baquerizo, M., 2019. Obscure soil microbes and where to find them. ISME J 13, 2120–2124. 10.1038/s41396-019-0405-0.

Delgado-Baquerizo, M., Maestre, F.T., Reich, P.B., Jeffries, T.C., Gaitan, J.J., Encinar, D., Berdugo, M., Campbell, C.D., Singh, B.K., 2016. Microbial diversity drives multifunctionality in terrestrial ecosystems. Nat Commun 7. 10.1038/ncomms10541.

Dragone, N.B., Hoffert, M., Strickland, M.S., Fierer, N., 2024. Taxonomic and genomic attributes of oligotrophic soil bacteria. ISME Commun 4, ycae081. 10.1093/ismeco/ycae081.

Duşa, A., 2024. venn: Draw Venn Diagrams. https://CRAN.R-project.org/package=venn.

Feller, F.M., Richtsmeier, P., Wege, M., Philipp, B., 2021. Comparative Analysis of Bile-Salt Degradation in Sphingobium sp. Strain Chol11 and Pseudomonas stutzeri Strain Chol1 Reveals Functional Diversity of Proteobacterial Steroid Degradation Enzymes and Suggests a Novel Pathway for Side Chain Degradation. Appl Environ Microbiol 87, e0145321. 10.1128/aem.01453-21.

Graves, S., Piepho, H.-P., Selzer, M.L., 2015. Package ‘multcompView’. Visualizations of paired comparisons.

Green, S.J., Prakash, O., Jasrotia, P., Overholt, W.A., Cardenas, E., Hubbard, D., Tiedje, J.M., Watson, D.B., Schadt, C.W., Brooks, S.C., Kostka, J.E., 2012. Denitrifying bacteria from the genus Rhodanobacter dominate bacterial communities in the highly contaminated subsurface of a nuclear legacy waste site. Appl Environ Microbiol 78, 1039–1047. 10.1128/aem.06435-11.

Grzyb, A., Wolna-Maruwka, A., Łukowiak, R., Ceglarek, J., 2022. Spatial and Temporal Variability of the Microbiological and Chemical Properties of Soils under Wheat and Oilseed Rape Cultivation. Agronomy 12, 2259.

Hinsu, A., Dumadiya, A., Joshi, A., Kotadiya, R., Andharia, K., Koringa, P., Kothari, R., 2021. To culture or not to culture: a snapshot of culture-dependent and culture-independent bacterial diversity from peanut rhizosphere. PeerJ 9, e12035. 10.7717/peerj.12035.

Hothorn, T., Bretz, F., Westfall, P., Heiberger, R.M., Schuetzenmeister, A., Scheibe, S., Hothorn, M.T., 2016. Package ‘multcomp’. Simultaneous inference in general parametric models. Project for Statistical Computing, Vienna, Austria.

Ivanova, A.A., Kulichevskaya, I.S., Merkel, A.Y., Toshchakov, S.V., Dedysh, S.N., 2016. High Diversity of Planctomycetes in Soils of Two Lichen-Dominated Sub-Arctic Ecosystems of Northwestern Siberia. Front Microbiol Volume 7 - 2016. 10.3389/fmicb.2016.02065.

Kielak, A.M., Barreto, C.C., Kowalchuk, G.A., van Veen, J.A., Kuramae, E.E., 2016. The Ecology of Acidobacteria: Moving beyond Genes and Genomes. Front Microbiol Volume 7 - 2016. 10.3389/fmicb.2016.00744.

Köninger, J., Panagos, P., Jones, A., Briones, M.J.I., Orgiazzi, A., 2022. In defence of soil biodiversity: Towards an inclusive protection in the European Union. Biological Conservation 268, 109475. 10.1016/j.biocon.2022.109475.

Kurm, V., Van Der Putten, W.H., Hol, W.H.G., 2019a. Cultivation-success of rare soil bacteria is not influenced by incubation time and growth medium. PLoS One 14. 10.1371/journal.pone.0210073.

Kurm, V., van der Putten, W.H., Weidner, S., Geisen, S., Snoek, B.L., Bakx, T., Hol, W.H.G., 2019b. Competition and predation as possible causes of bacterial rarity. Environ Microbiol 21, 1356–1368. 10.1111/1462-2920.14569.

Lee, S.A., Park, J., Chu, B., Kim, J.M., Joa, J.H., Sang, M.K., Song, J., Weon, H.Y., 2016. Comparative analysis of bacterial diversity in the rhizosphere of tomato by culture-dependent and -independent approaches. J Microbiol 54, 823–831. 10.1007/s12275-016-6410-3.

Lennon, J.T., Jones, S.E., 2011. Microbial seed banks: the ecological and evolutionary implications of dormancy. Nat Rev Micro 9, 119–130. http://www.nature.com/nrmicro/journal/v9/n2/suppinfo/nrmicro2504_S1.html.

Lewis, W.H., Tahon, G., Geesink, P., Sousa, D.Z., Ettema, T.J.G., 2020. Innovations to culturing the uncultured microbial majority. Nature Reviews Microbiology 10.1038/s41579-020-00458-8.

Liang, R., Lau, M., Vishnivetskaya, T., Lloyd Karen, G., Wang, W., Wiggins, J., Miller, J., Pfiffner, S., Rivkina Elizaveta, M., Onstott Tullis, C., 2019. Predominance of Anaerobic, Spore-Forming Bacteria in Metabolically Active Microbial Communities from Ancient Siberian Permafrost. Appl Environ Microbiol 85, e00560–00519. 10.1128/AEM.00560-19.

Lynch, M.D.J., Neufeld, J.D., 2015. Ecology and exploration of the rare biosphere. Nature Reviews Microbiology 13, 217–229 10.1038/nrmicro3400.

Mandakovic, D., Maldonado, J., Pulgar, R., Cabrera, P., Gaete, A., Urtuvia, V., Seeger, M., Cambiazo, V., González, M., 2018. Microbiome analysis and bacterial isolation from Lejía Lake soil in Atacama Desert. Extremophiles 22, 665–673 10.1007/s00792-018-1027-6.

Mitra, M., Nguyen, K.M., Box, T.W., Gilpin, J.S., Hamby, S.R., Berry, T.L., Duckett, E.H., 2020. Isolation and characterization of a novel Sphingobium yanoikuyae strain variant that uses biohazardous saturated hydrocarbons and aromatic compounds as sole carbon sources. F1000Res 9, 767. 10.12688/f1000research.25284.1.

Molina-Menor, E., Gimeno-Valero, H., Pascual, J., Peretó, J., Porcar, M., 2021. High Culturable Bacterial Diversity From a European Desert: The Tabernas Desert. Front Microbiol 11. 10.3389/fmicb.2020.583120.

Nguyen, T.M., Seo, C., Ji, M., Paik, M.-J., Myung, S.-W., Kim, J., 2018. Effective Soil Extraction Method for Cultivating Previously Uncultured Soil Bacteria. Appl Environ Microbiol 84, e01145–01118. 10.1128/aem.01145-18.

Nixon, S.L., Daly, R.A., Borton, M.A., Solden, L.M., Welch, S.A., Cole, D.R., Mouser, P.J., Wilkins, M.J., Wrighton, K.C., 2019. Genome-Resolved Metagenomics Extends the Environmental Distribution of the <i>Verrucomicrobia</i> Phylum to the Deep Terrestrial Subsurface. mSphere 4, 10.1128/msphere.00613-00619. doi:10.1128/msphere.00613-19.

Pascoal, F., Branco, P., Torgo, L., Costa, R., Magalhães, C., 2025. Definition of the microbial rare biosphere through unsupervised machine learning. Communications Biology 8, 544. 10.1038/s42003-025-07912-4.

Pascoal, F., Costa, R., Magalhães, C., 2021. The microbial rare biosphere: current concepts, methods and ecological principles. FEMS Microbiol Ecol 97, fiaa227.

Pascoal, F., Magalhães, C., Costa, R., 2020. The Link Between the Ecology of the Prokaryotic Rare Biosphere and Its Biotechnological Potential. Front Microbiol Volume 11 - 2020. 10.3389/fmicb.2020.00231.

Pascual, J., Blanco, S., García-López, M., García-Salamanca, A., Bursakov, S.A., Genilloud, O., Bills, G.F., Ramos, J.L., van Dillewijn, P., 2016. Assessing Bacterial Diversity in the Rhizosphere of <italic>Thymus zygis</italic> Growing in the Sierra Nevada National Park (Spain) through Culture-Dependent and Independent Approaches. PLoS One 11, e0146558. 10.1371/journal.pone.0146558.

Pédron, J., Guyon, L., Lecomte, A., Blottière, L., Chandeysson, C., Rochelle-Newall, E., Raynaud, X., Berge, O., Barny, M.A., 2020. Comparison of Environmental and Culture-Derived Bacterial Communities through 16S Metabarcoding: A Powerful Tool to Assess Media Selectivity and Detect Rare Taxa. Microorganisms 8. 10.3390/microorganisms8081129.

Pham, V.H., Kim, J., 2012. Cultivation of unculturable soil bacteria. Trends Biotechnol 30, 475–484. 10.1016/j.tibtech.2012.05.007.

Pulido-Chavez, M.F., Randolph, J.W.J., Zalman, C., Larios, L., Homyak, P.M., Glassman, S.I., 2023. Rapid bacterial and fungal successional dynamics in first year after chaparral wildfire. Molecular ecology 32, 1685–1707. 10.1111/mec.16835.

Rachid, C.T., Santos, A.L., Piccolo, M.C., Balieiro, F.C., Coutinho, H.L., Peixoto, R.S., Tiedje, J.M., Rosado, A.S., 2013. Effect of sugarcane burning or green harvest methods on the Brazilian Cerrado soil bacterial community structure. PLoS One 8, e59342. 10.1371/journal.pone.0059342.

Riddley, M., Hepp, S., Hardeep, F.N.U., Nayak, A., Liu, M., Xing, X., Zhang, H., Liao, J., 2025. Differential roles of deterministic and stochastic processes in structuring soil bacterial ecotypes across terrestrial ecosystems. Nature Communications 16, 2337. 10.1038/s41467-025-57526-x.

Schloss, P.D., Westcott, S.L., Ryabin, T., Hall, J.R., Hartmann, M., Hollister, E.B., Lesniewski, R.A., Oakley, B.B., Parks, D.H., Robinson, C.J., Sahl, J.W., Stres, B., Thallinger, G.G., Van Horn, D.J., Weber, C.F., 2009. Introducing mothur: Open-source, platform-independent, community-supported software for describing and comparing microbial communities. Appl Environ Microbiol 75, 7537–7541.

Shade, A., Hogan, C.S., Klimowicz, A.K., Linske, M., McManus, P.S., Handelsman, J., 2012. Culturing captures members of the soil rare biosphere. Environ Microbiol 14, 2247–2252. 10.1111/j.1462-2920.2012.02817.x.

Siles, J.A., Hendrickson, A.J., Terry, N., 2022. Coupling of metataxonomics and culturing improves bacterial diversity characterization and identifies a novel Rhizorhapis sp. with metal resistance potential in a multi-contaminated waste sediment. J Environ Manage 322, 116132. 10.1016/j.jenvman.2022.116132.

Steen, A.D., Crits-Christoph, A., Carini, P., DeAngelis, K.M., Fierer, N., Lloyd, K.G., Cameron Thrash, J., 2019. High proportions of bacteria and archaea across most biomes remain uncultured. ISME J. 10.1038/s41396-019-0484-y.

Stefani, F.O.P., Bell, T.H., Marchand, C., de la Providencia, I.E., El Yassimi, A., St-Arnaud, M., Hijri, M., 2015. Culture-Dependent and -Independent Methods Capture Different Microbial Community Fractions in Hydrocarbon-Contaminated Soils. PLoS One 10, e0128272. 10.1371/journal.pone.0128272.

Stewart, K.A., Setlow, P., 2013. Numbers of individual nutrient germinant receptors and other germination proteins in spores of Bacillus subtilis. J Bacteriol 195, 3575–3582. 10.1128/jb.00377-13.

Thompson, L.R., Sanders, J.G., McDonald, D., Amir, A., Ladau, J., Locey, K.J., Prill, R.J., Tripathi, A., Gibbons, S.M., Ackermann, G., Navas-Molina, J.A., Janssen, S., Kopylova, E., Vázquez-Baeza, Y., González, A., et al., 2017. A communal catalogue reveals Earth’s multiscale microbial diversity. Nature 551, 457–463. 10.1038/nature24621.

Thrash, J.C., 2019. Culturing the Uncultured: Risk versus Reward. mSystems 4, e00130–00119. doi:10.1128/mSystems.00130-19.

Thrash, J.C., 2021. Towards culturing the microbe of your choice. Environmental Microbiology Reports 13, 36–41. 10.1111/1758-2229.12898.

VanInsberghe, D., Hartmann, M., Stewart, G.R., Mohn, W.W., 2013. Isolation of a Substantial Proportion of Forest Soil Bacterial Communities Detected via Pyrotag Sequencing. Appl Environ Microbiol 79, 2096–2098. 10.1128/aem.03112-12.

Wang, M., Noor, S., Huan, R., Liu, C., Li, J., Shi, Q., Zhang, Y.J., Wu, C., He, H., 2020. Comparison of the diversity of cultured and total bacterial communities in marine sediment using culture-dependent and sequencing methods. PeerJ 8, e10060. 10.7717/peerj.10060.

Wickham, H., 2016. ggplot2: Elegant Graphics for Data Analysis. Springer-Verlag New York.

Xu, Y., Vetsigian, K., 2017. Phenotypic variability and community interactions of germinating Streptomyces spores. Sci Rep 7, 699. 10.1038/s41598-017-00792-7.

Youseif, S.H., Abd El-Megeed, F.H., Humm, E.A., Maymon, M., Mohamed, A.H., Saleh, S.A., Hirsch, A.M., 2021. Comparative Analysis of the Cultured and Total Bacterial Community in the Wheat Rhizosphere Microbiome Using Culture-Dependent and Culture-Independent Approaches. Microbiol Spectr 9, e0067821. 10.1128/Spectrum.00678-21.

Zhang, C., Liu, S., Guo, Q., Li, D., Li, Z., Ma, Q., Liu, H., Zhao, Q., Liu, H., Ding, Z., Gong, W., Gao, Y., 2024. Sphingobium sp. V4, a bacterium degrading multiple allelochemical phenolic acids. Annals of Microbiology 74, 5. 10.1186/s13213-024-01750-1.

