## Supplementary Material for "Accessing rare bacterial biosphere of soil through culturing: a comparative study of culture media effectiveness integrated with metataxonomics"

*for*

**The supplementary material includes Figures S1-S4 and Tables S1-S12**

Fig. S1. Rank-abundance curves of bacterial culture-independent communities in the three studied soils. The x-axis represents the rank order of OTUs (from most to least abundant), while the y-axis indicates their abundance on a logarithmic scale. Rarity thresholds of 0.1% and 0.01% are indicated.

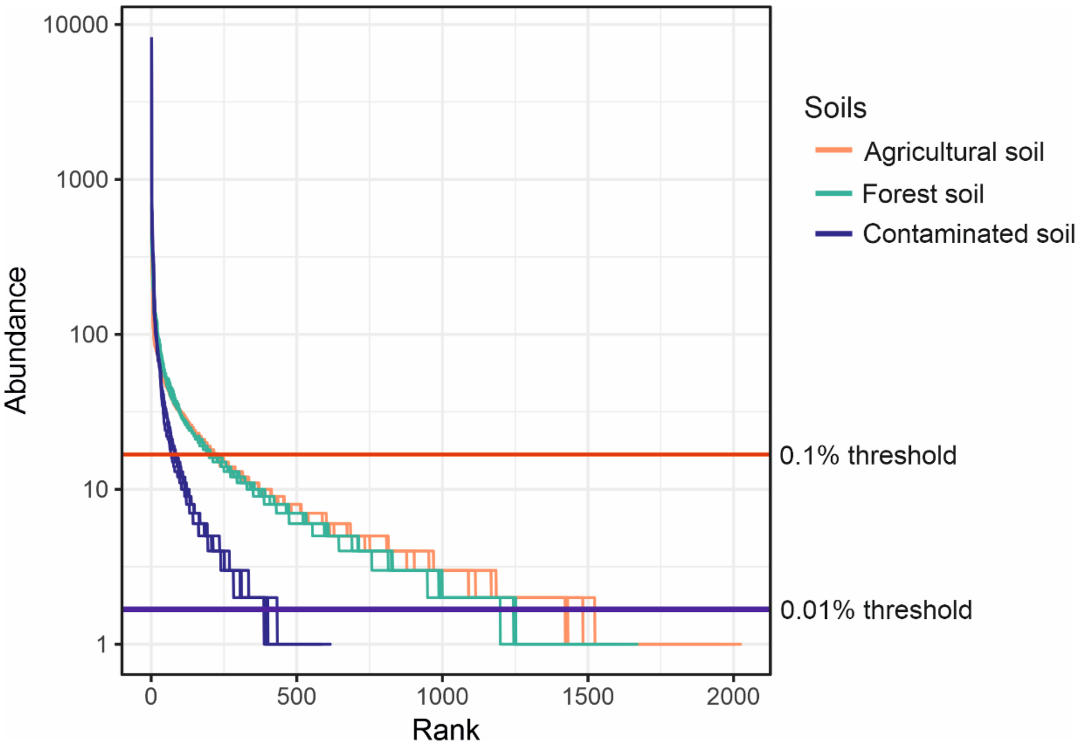

Fig. S2. Non-metric multidimensional scaling (NMDS) ordination of samples according to bacterial community structure, based on the culture-independent approach and the culture-dependent approach using TSA (tryptic soy agar), SEM (soil extract medium), R2A (Reasoner's 2A), and 1:20 R2A (Reasoner's 2A diluted 20-fold) from agricultural, forest, and contaminated soils. NMDS analysis was done at OTU level taking advantage of the *metaMDS* function in the R package vegan ver. 2.6–4. The significance of the differences shown by the NMDS ordination was statistically tested using permutational analysis of variance (PERMANOVA) based on Bray-Curtis dissimilarities with 9,999 permutations, implemented in the R package vegan

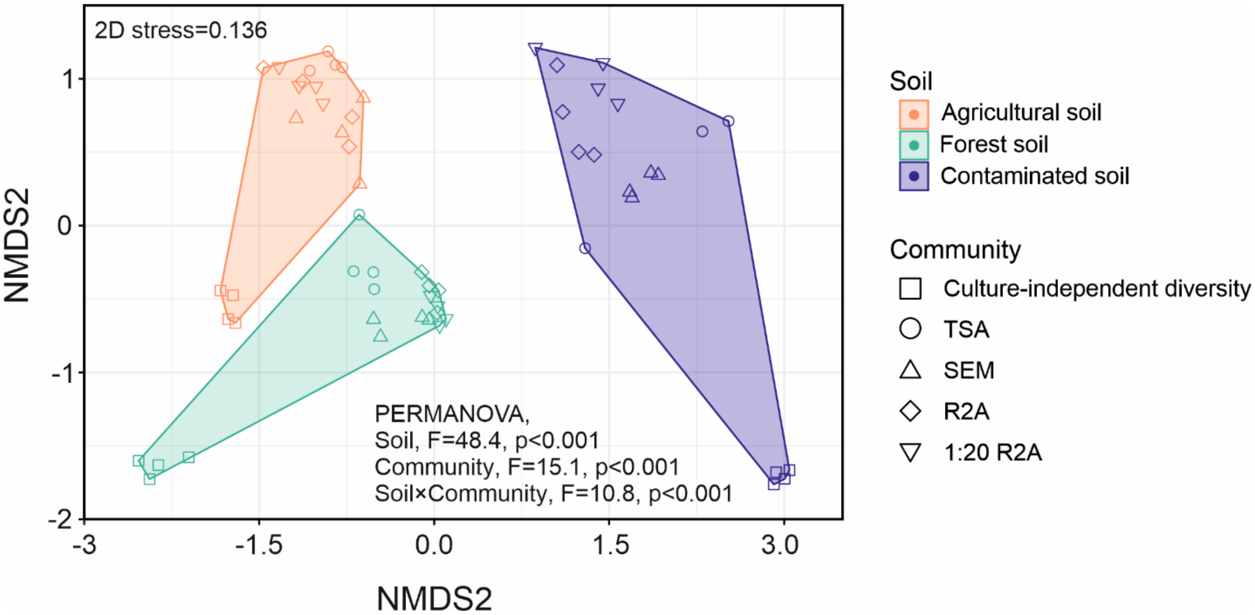

Fig. S3. Taxonomic composition at class level of the bacterial culture-independent community in the agricultural, forest and contaminated soils.

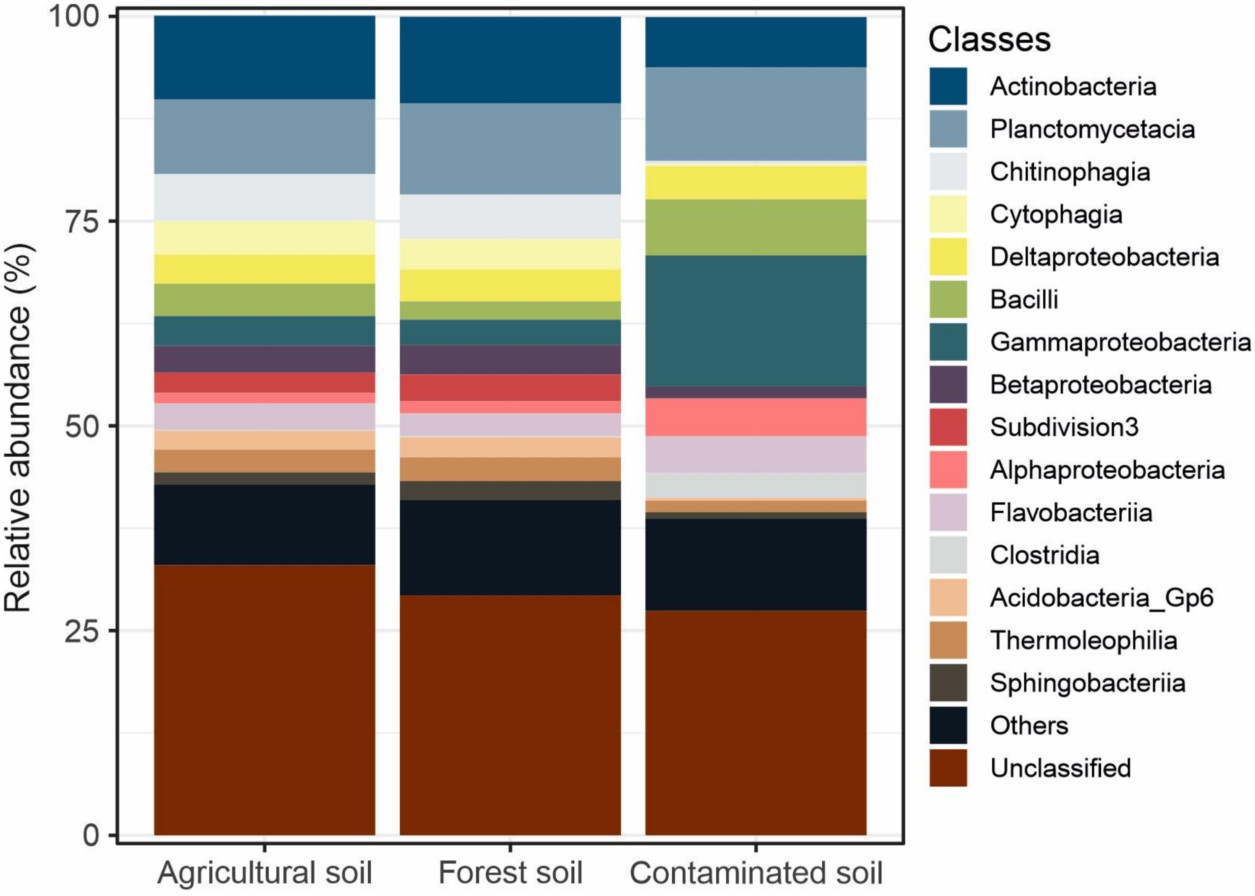

Fig. S4. Rarefaction curves of bacterial culture-independent communities in the agricultural, forests and contaminated soils.

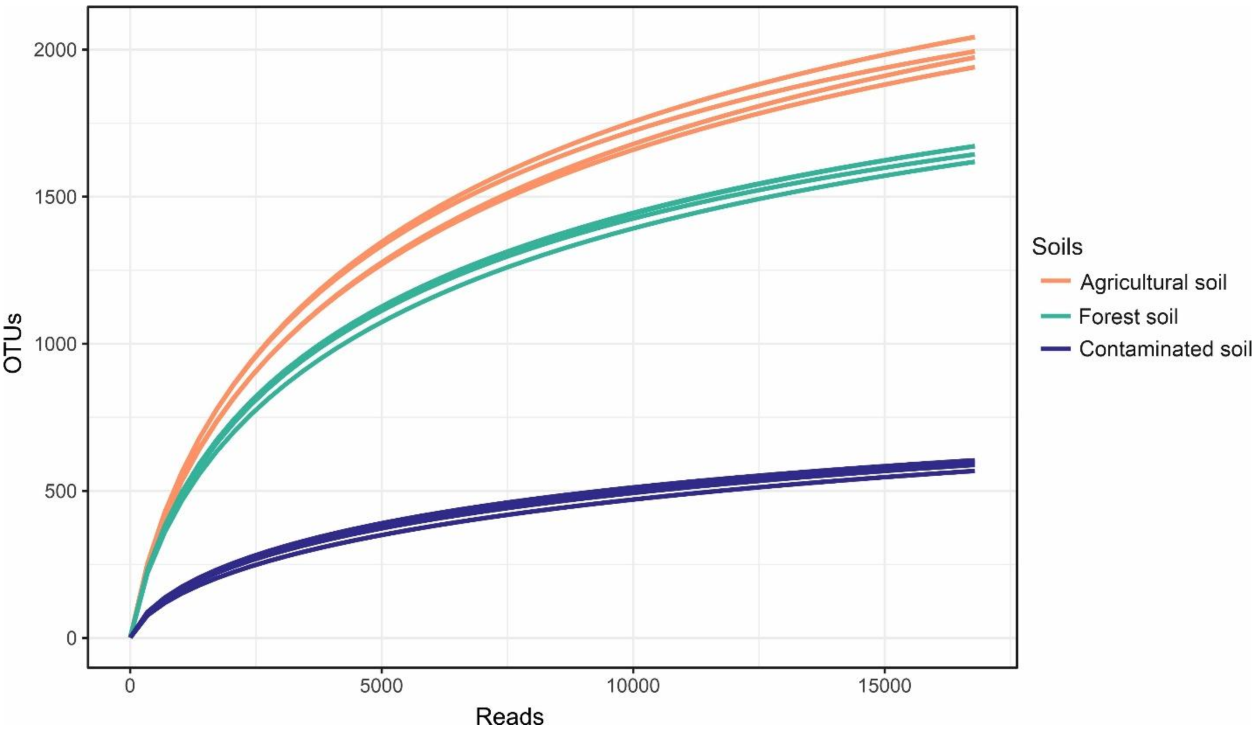

Table S1. Main properties of the three soils used in the study. Values represent means ± standard deviations (n=3). EC, electrical conductivity. TC, total carbon. TOC, total organic carbon. TN, total nitrogen. TP, total phosphorus. C:N, TOC:TN. Values represent means ± standard deviations.

| Property | Agricultural soil | Forest soil | Contaminated soil |
|---|---|---|---|
| Sand (%) | 42.0 ± 0.8 | 30.3 ± 1.0 | 38.7 ± 0.6 |
| Silt (%) | 29.3 ± 0.5 | 42.8 ± 1.3 | 21.7 ± 0.6 |
| Clay (%) | 28.8 ± 0.1 | 27.0 ± 1.2 | 39.7 ± 0.6 |
| Texture | Clay Loam | Clay | Loam |
| pH | 7.0 ± 0.1 | 6.5 ± 0.1 | 8.2 ± 0.1 |
| EC (mS $cm^{-1}$) | 0.185 ± 0.008 | 0.310 ± 0.002 | 1.19 ± 0.06 |
| TC (mg $g^{-1}$) | 25.2 ± 3.8 | 51.7 ± 4.3 | 534 ± 14 |
| TOC (mg $g^{-1}$) | 24.9 ± 2.9 | 51.0 ± 5.3 | 533 ± 7 |
| TN (mg $g^{-1}$) | 2.2 ± 0.5 | 2.7 ± 0.1 | 2.7 ± 0.3 |
| TP (mg $g^{-1}$) | 0.88 ± 0.05 | 0.44 ± 0.01 | 0.69 ± 0.19 |
| C:N | 11.7 ± 1.2 | 19.1 ± 1.2 | 195 ± 20 |
| Bacterial abundance (16S rRNA gene copy number $g^{-1}$ soil) | $1.95 \times 10^{10} \pm 2.28 \times 10^{9}$ | $9.85 \times 10^{10} \pm 3.1 \times 10^{9}$ | $5.32 \times 10^{8} \pm 1.95 \times 10^{7}$ |

Table S2. Nutrient composition of the culture media used for the culture-dependent approach. Values represent means ± standard deviations (n=3). TSA, tryptic soy agar. R2A, Reasoner's 2A. 1:20 R2A, Reasoner's 2A diluted 20-fold. SEM, soil extracted medium using agricultural, forest, and contaminated soil. TOC, total organic carbon. N, total nitrogen. P, total phosphorus. K, total potassium. Ca, total calcium. Mg, total magnesium. Na, total sodium. Values represent means ± standard deviations.

| Nutrient | TSA | R2A | 1:20 R2A | SEM agricultural soil | SEM forest soil | SEM contaminated soil |
|---|---|---|---|---|---|---|
| TOC (g $l^{-1}$) | 9.55 ± 0.08 | 1.12 ± 0.12 | 0.041 ± 0.081 | 0.166 ± 0.034 | 0.149 ± 0.012 | 0.216 ± 0.064 |
| N (mg $l^{-1}$) | 2140 ± 40 | 154.2 ± 6.5 | 7.23 ± 0.23 | 20.8 ± 1.23 | 12.2 ± 2.53 | 28.8 ± 3.45 |
| P (mg $l^{-1}$) | 5389 ± 53 | 107.9 ± 8.1 | 5.41 ± 1.25 | 8.16 ± 0.98 | 0.731 ± 0.045 | 3.64 ± 0.56 |
| K (mg $l^{-1}$) | 1370 ± 23 | 164.1 ± 12.1 | 8.90 ± 0.67 | 17.7 ± 1.11 | 8.80 ± 0.99 | 23.3 ± 1.2 |
| Ca (mg $l^{-1}$) | 7.20 ± 0.95 | 4.50 ± 0.65 | 0.612 ± 0.098 | 22.3 ± 0.98 | 12.6 ± 0.65 | 44.7 ± 1.4 |
| Mg (mg $l^{-1}$) | 11.9 ± 1.1 | 6.91 ± 0.99 | 0.405 ± 0.087 | 6.90 ± 0.40 | 6.50 ± 0.43 | 31.0 ± 1.21 |
| Na (mg $l^{-1}$) | 2288 ± 92 | 313 ± 21 | 15.9 ± 0.76 | 6.70 ± 0.56 | 6.10 ± 0.96 | 393 ± 11 |

Table S3. Taxonomic composition at genus level of the bacterial culture-independent community and culture-dependent community using TSA (tryptic soy agar), SEM (soil extract medium), R2A (Reasoner's 2A), and 1:20 R2A (Reasoner's 2A diluted 20-fold) from agricultural soil. Values represent means ± standard deviations.

| Genus | Culture-independent | TSA | SEM | R2A | 1:20 R2A |
|---|---|---|---|---|---|
| *Bacillus* | 2.9±1.14 | 18.75±4.51 | 7.74±1.53 | 26.13±7.86 | 35.48±8.3 |
| *Stenotrophomonas* | 0.51±0.28 | 12.99±6.77 | 8.83±2.85 | 7.55±3.57 | 13.04±4.14 |
| *Pseudomonas* | 1.47±0.52 | 6.55±1.8 | 10.06±11.63 | 6.99±7.6 | 4.29±4.76 |
| *Nocardioides* | 1.02±0.29 | 0.04±0.01 | 12.96±2.39 | 1.42±1.04 | 4.22±2.38 |
| *Peribacillus* | 0.46±0.06 | 7±5.04 | 3.96±2.25 | 1.53±1.86 | 0.63±0.18 |
| *Variovorax* | 0.41±0.07 | 0.05±0.01 | 6.86±4.57 | 1.47±1.3 | 3.98±2.34 |
| *Flavobacterium* | 2.58±0.63 | 0.07±0.05 | 1.98±2.07 | 0.56±0.43 | 6.98±12.77 |
| *Chitinophaga* | 1.28±1.12 | 0.05±0.01 | 6.47±4.58 | 1.31±0.92 | 2.05±1.96 |
| *Neobacillus* | 2±0.53 | 0.05±0.01 | 4.57±2.7 | 0.36±0.53 | 0.46±0.31 |
| *Pedobacter* | 0.72±0.15 | 0.02±0.02 | 2.44±2.71 | 2.49±2.87 | 0.86±1.17 |
| *Achromobacter* | 0.17±0.06 | 2.97±2.14 | 0.55±0.32 | 2.08±1.61 | 0.25±0.31 |
| *Buttiauxella* | 0.09±0.03 | 1.53±1.02 | 0.5±0.43 | 2.1±1.29 | 1.71±1.03 |
| *Ramlibacter* | 0.77±0.11 | 0.02±0.01 | 4.45±4.8 | 0.03±0.02 | 0.59±0.49 |
| *Sphingobacterium* | 0.64±1.05 | 2±2.12 | 0.36±0.49 | 2.23±3.4 | 0.21±0.18 |
| *Chryseobacterium* | 1.01±0.77 | 2.21±3.9 | 0.11±0.07 | 0.43±0.23 | 0.29±0.37 |
| Gp6 | 3.68±0.47 | 0.02±0.01 | 0.03±0.01 | 0.02±0.01 | 0.02±0 |
| *Flavisolibacter* | 3.66±0.17 | | 0.06±0.06 | 0.01±0 | |
| *Streptomyces* | 0.69±0.1 | 0.39±0.38 | 1.3±0.68 | 0.21±0.21 | 0.67±0.5 |
| Gp4 | 3.17±0.46 | 0.01±0.01 | 0.01±0 | 0.02±0.01 | 0.01±0.01 |
| Spartobacteria g i s | 1.89±0.11 | 0.01±0 | 1±0.96 | 0.01±0.01 | 0.02±0.01 |
| *Paenibacillus* | 0.13±0.05 | 1.44±1.63 | 0.2±0.22 | 0.96±0.84 | 0.13±0.09 |
| *Lysobacter* | 0.86±0.01 | 0.02±0.01 | 0.4±0.21 | 1.35±1.77 | 0.09±0.07 |
| *Luteibacter* | 0.02±0.01 | 0.01±0.01 | 0.36±0.39 | 0.75±0.78 | 1.04±0.89 |
| *Psychrobacillus* | 0.03±0.02 | 2.06±4.02 | 0.01±0.01 | 0.02±0.02 | 0.02±0.01 |
| *Paraburkholderia* | 0.07±0.1 | 0.04±0.05 | 0.08±0.08 | 0.01±0.01 | 1.93±3.78 |
| *Pseudoxanthomonas* | 0.03±0.02 | 0.04±0.07 | 0.61±0.54 | 0.69±1.18 | 0.76±1.4 |
| Subdivision3 g i s | 2.09±0.29 | | 0.01±0.01 | | 0.01±0.01 |
| *Olivibacter* | 0.04±0.07 | 0.01±0.01 | 1.51±2.08 | 0.12±0.13 | 0.17±0.34 |
| *Arthrobacter* | 0.28±0.07 | 0.14±0.16 | 0.69±0.45 | 0.47±0.24 | 0.25±0.06 |
| *Microbacterium* | 0.18±0.05 | 0.26±0.25 | 0.35±0.19 | 0.5±0.54 | 0.51±0.38 |
| *Skermanella* | 1.51±0.18 | | | 0.01±0.01 | |
| *Microlunatus* | 1.37±0.24 | | 0.01±0 | 0.01±0.01 | |
| *Massilia* | 1.23±0.21 | 0.01±0 | 0.07±0.09 | 0.01±0.01 | 0.06±0.04 |
| *Adhaeribacter* | 1.32±0.17 | | | | |
| *Agromyces* | 0.25±0.09 | 0.06±0.08 | 0.32±0.14 | 0.32±0.33 | 0.37±0.22 |
| *Chryseolinea* | 1.23±0.46 | 0.01±0 | 0.02±0.03 | 0.01±0.01 | 0.01±0.01 |
| *Rhodococcus* | 0.04±0.02 | 0.03±0.02 | 0.42±0.3 | 0.16±0.17 | 0.37±0.6 |
| *Serratia* | 0.08±0.03 | 0.26±0.05 | 0.16±0.06 | 0.3±0.05 | 0.17±0.03 |
| *Rubrobacter* | 0.96±0.31 | | | | |
| WPS-1 g i s | 0.87±0.17 | | | | 0.01±0.01 |
| *Pantoea* | 0.08±0.05 | 0.67±0.76 | 0.01±0.02 | 0.03±0.03 | 0.03±0.05 |
| *Mycobacterium* | 0.32±0.03 | | 0.16±0.14 | 0.02±0 | 0.3±0.27 |
| *Xanthomonas* | 0.02±0.01 | 0.03±0.03 | 0.36±0.48 | 0.09±0.13 | 0.23±0.12 |
| *Ferruginibacter* | 0.7±0.25 | | 0.01±0.02 | | |
| *Cellvibrio* | 0.69±0.67 | | | | |
| *Mucilaginibacter* | 0.45±0.22 | | 0.11±0.11 | 0.12±0.23 | 0.01±0.01 |
| *Dyadobacter* | 0.23±0.08 | | 0.4±0.3 | 0.02±0.04 | 0.02±0.02 |
| Others | 11.45±0.59 | 2.01±1.33 | 2.63±0.71 | | 1.44±0.75 |
| Unassigned | 44.38±1.23 | 38.15±9.56 | 16.85±7.92 | 36.13±13.27 | 16.3±5.39 |

Table S4. Taxonomic composition at genus level of the bacterial culture-independent community and culture-dependent community using TSA (tryptic soy agar), SEM (soil extract medium), R2A (Reasoner's 2A), and 1:20 R2A (Reasoner's 2A diluted 20-fold) from forest soil. Values represent means ± standard deviations in percentages.

| Genus | Culture-independent | TSA | SEM | R2A | 1:20 R2A |
|---|---|---|---|---|---|
| *Pseudomonas* | 0.41±0.1 | 21.05±8.01 | 50.4±2.32 | 32.83±4.35 | 51.3±7.25 |
| *Serratia* | 0.09±0.03 | 27.39±20.77 | 1.35±1.11 | 19.85±9.72 | 5.45±4.95 |
| *Chitinophaga* | 0.11±0.04 | 0.04±0.02 | 9.73±9.5 | 5.43±10.44 | 15.89±9.26 |
| *Peribacillus* | 0.18±0.07 | 23.02±25.19 | 2.43±1.64 | 5.1±5.77 | 0.08±0.06 |
| *Pedobacter* | 0.11±0.12 | 0.03±0.01 | 6.07±2.78 | 10.58±7.52 | 7.41±6.8 |
| *Flavobacterium* | 3.6±2.81 | 0.75±0.56 | 8.06±3.28 | 8.73±4.4 | 4.6±3.15 |
| *Buttiauxella* | 0.03±0.02 | 6.57±6.84 | 0.72±0.87 | 5.31±3.86 | 1.78±2.05 |
| *Stenotrophomonas* | 0.13±0.02 | 2.72±1.78 | 0.46±0.38 | 1.38±1.74 | 1±1.11 |
| *Phyllobacterium* | 0.01±0 | 0.01±0.01 | 2.79±3.34 | 1.64±1.86 | 0.82±0.82 |
| *Variovorax* | 0.14±0.03 | 0.06±0.02 | 2.33±3.05 | 0.42±0.36 | 2.27±1.97 |
| *Aeromonas* | 0.03±0.03 | 2.51±4.72 | 0.96±1.78 | 0.72±1.39 | 0.51±0.99 |
| *Neobacillus* | 0.28±0.12 | 0.04±0.03 | 4.06±1.31 | 0.12±0.14 | 0.28±0.19 |
| *Sporosarcina* | 0.02±0.01 | 4.44±5.85 | | | |
| *Rhodococcus* | 0.03±0.02 | 0.18±0.13 | 0.45±0.39 | 0.26±0.16 | 2.4±2.09 |
| *Bacillus* | 0.2±0.02 | 0.64±0.17 | 1.68±2.8 | 0.35±0.25 | 0.22±0.13 |
| *Sphingobacterium* | 0.03±0.01 | 2.37±4.16 | 0.02±0.02 | 0.11±0.13 | |
| *Psychrobacillus* | 0.02±0.01 | 1.6±2.76 | | | |
| *Agromyces* | 0.04±0.02 | 0.11±0.1 | 0.11±0.06 | 0.99±0.83 | 0.22±0.33 |
| *Nocardioides* | 0.56±0.07 | 0.05±0.01 | 0.5±0.64 | 0.18±0.12 | 0.64±0.46 |
| *Streptomyces* | 0.19±0.05 | 0.05±0.03 | 0.36±0.2 | 0.17±0.01 | 0.61±0.23 |
| Spartobacteria g i s | 8.42±1.62 | 0.02±0.02 | 0.78±0.42 | | |
| *Lysobacter* | 0.13±0.07 | 0.01±0.01 | 0.25±0.21 | 0.21±0.07 | 0.17±0.05 |
| *Oceanobacillus* | | 0.57±1.04 | | | |
| *Paenibacillus* | 0.02±0.01 | 0.07±0.04 | 0.22±0.07 | 0.19±0.3 | 0.01±0.01 |
| *Achromobacter* | 0.03±0.01 | 0.44±0.84 | 0.02±0.02 | 0.02±0.04 | 0±0 |
| *Chryseobacterium* | 0.01±0.02 | 0.28±0.46 | 0.03±0.02 | 0.04±0.03 | 0.04±0.05 |
| *Staphylococcus* | 0.01±0.01 | 0.36±0.69 | | 0.01±0.01 | 0.01±0.01 |
| *Mucilaginibacter* | 0.06±0.02 | | 0.16±0.1 | 0.04±0.06 | 0.12±0.19 |
| *Paraburkholderia* | 0.01±0.01 | 0.01±0.01 | 0.13±0.04 | 0.01±0.01 | 0.14±0.16 |
| *Mycobacterium* | 0.13±0.07 | | 0.03±0.03 | 0.01±0.01 | 0.24±0.17 |
| *Mesobacillus* | 0±0 | | 0.07±0.08 | 0.11±0.02 | 0.1±0.02 |
| *Dyadobacter* | 0.02±0.02 | | 0.18±0.18 | 0.01±0.02 | 0.04±0.05 |
| *Ramlibacter* | 0.33±0.05 | 0.01±0.01 | 0.12±0.12 | | 0.02±0.02 |
| *Leifsonia* | | | 0.02±0.01 | 0.08±0.11 | 0.03±0.02 |
| *Fictibacillus* | | | 0.02±0.03 | 0.06±0.02 | 0.04±0.02 |
| *Nocardia* | | | 0.1±0.06 | | 0.01±0.02 |
| *Arthrobacter* | 0.02±0.01 | 0.04±0.05 | 0.01±0.01 | 0.01±0.01 | 0.05±0.04 |
| *Caballeronia* | 0.01±0.01 | | 0.04±0.02 | | 0.06±0.05 |
| *Rhizobium* | | | 0.01±0 | 0.06±0.08 | 0.02±0.01 |
| *Marmoricola* | 0.04±0.02 | | 0.05±0.05 | 0.01±0.01 | 0.04±0.03 |
| *Luteibacter* | 0.01±0 | 0.01±0.01 | 0.04±0.06 | | 0.03±0.03 |
| *Polaromonas* | | | 0.02±0.03 | 0.02±0.02 | 0.03±0.02 |
| *Edaphobacter* | 0.07±0.03 | | 0.05±0.05 | | 0.02±0.02 |
| *Acinetobacter* | | 0.02±0.04 | 0.01±0.01 | 0.03±0.05 | 0.01±0.01 |
| *Brucella* | | | 0.01±0.01 | 0.02±0.03 | 0.03±0.06 |
| *Roseimicrobium* | 0.01±0 | | 0.06±0.04 | | |
| *Singulisphaera* | 0.09±0.02 | | 0.04±0.02 | | |
| *Sediminibacterium* | 0.04±0.01 | | 0.05±0.04 | | |
| *Kitasatospora* | 0.01±0.01 | | 0±0.01 | 0.01±0.01 | 0.03±0.03 |
| *Kangiella* | | | 0.01±0.01 | 0.01±0.01 | 0.02±0.01 |
| *Pseudoxanthomonas* | 0.01±0.01 | 0.01±0 | 0.02±0.02 | | 0.01±0 |
| *Mesorhizobium* | | | 0.02±0.01 | | 0.02±0.01 |
| Gp6 | 10.48±0.91 | 0.02±0.01 | 0.02±0.01 | | |
| Others | 24.27±1.84 | 0.14±0.03 | 0.35±0.16 | 0.14±0.08 | 0.21±0.08 |
| (Unassigned) | 49.58±1.94 | 4.34±4.35 | 4.54±1.97 | 4.68±2.64 | 2.97±1.8 |

Table S5. Taxonomic composition at genus level of the bacterial culture-independent community and culture-dependent community using TSA (tryptic soy agar), SEM (soil extract medium), R2A (Reasoner's 2A), and 1:20 R2A (Reasoner's 2A diluted 20-fold) from contaminated soil. Values represent means ± standard deviations. Values represent means ± standard deviations in percentages.

| Genus | Culture -independent | TSA | SEM | R2A | 1:20 R2A |
|---|---|---|---|---|---|
| *Pseudomonas* | 0.54±0.1 | 6.87±9 | 34±11.95 | 32.48±14.32 | 14.51±12.85 |
| *Streptomyces* | 0.21±0.05 | 1.62±0.2 | 2.49±2.02 | 28.88±8.51 | 21.15±9.71 |
| *Lysobacter* | 0.53±0.16 | 46.83±6.99 | 13.85±3.53 | 0.45±0.43 | 0.13±0.04 |
| *Mesobacillus* | 0.41±0.06 | 3.85±0.95 | 25.38±9.26 | 1.28±0.9 | 13.29±9.77 |
| *Bacillus* | 0.12±0.03 | 16.63±4.19 | 0.14±0.04 | 21.63±7.36 | 7.03±1.47 |
| *Nocardioides* | 0.18±0.05 | 0.09±0.02 | 4.33±3.43 | 2.68±1.22 | 12.71±10.06 |
| *Fictibacillus* | 0.04±0.02 | 0.08±0.02 | 0.06±0.03 | 2.12±1.16 | 14.85±11.78 |
| *Rhodococcus* | 0.03±0.03 | 0.09±0.04 | 0.21±0.31 | 2.38±2.52 | 7.07±12.22 |
| *Mycobacterium* | 0.06±0.01 | 0.02±0.01 | 3.6±4.05 | 0.03±0.01 | 1.18±0.53 |
| *Solibacillus* | 0.01±0.01 | 2.86±1.8 | | | 0.01±0.01 |
| *Ornithinimicrobium* | 0.03±0.01 | 0.52±0.36 | 0.56±0.16 | 0.26±0.1 | 0.07±0.03 |
| *Peribacillus* | 0.02±0.01 | 0.14±0.12 | 0.16±0.23 | 0.24±0.32 | 0.68±0.75 |
| *Neobacillus* | 0.01±0.01 | 0.01±0 | 0.03±0.01 | 0.16±0.12 | 0.89±1.29 |
| *Janibacter* | | 0.29±0.42 | 0.33±0.29 | 0.17±0.32 | |
| *Paenibacillus* | | 0.01±0.02 | 0.01±0 | 0.63±1.2 | 0.02±0.02 |
| *Virgibacillus* | | 0.65±1.12 | | | |
| *Alkalihalobacillus* | | | 0.05±0.04 | 0.11±0.07 | 0.31±0.41 |
| *Pusillimonas* | 0.16±0.07 | | 0.44±0.21 | 0.01±0.01 | |
| *Pseudactinotalea* | | | 0.24±0.11 | 0.07±0.04 | 0.11±0.11 |
| *Dietzia* | | 0.04±0.01 | 0.28±0.53 | | 0.04±0.07 |
| *Serratia* | 0.08±0.01 | 0.1±0.01 | 0.11±0 | 0.08±0.03 | 0.07±0.02 |
| *Tsukamurella* | | | | 0.29±0.33 | |
| *Georgenia* | | 0.01±0.01 | 0.21±0.13 | 0.03±0.02 | 0.04±0.03 |
| *Chitinophaga* | 0.05±0.02 | 0.08±0.02 | 0.07±0.01 | 0.06±0.01 | 0.04±0.01 |
| *Cellulosimicrobium* | | 0.01±0.01 | | | 0.22±0.45 |
| *Flavobacterium* | 0.07±0.04 | 0.05±0.01 | 0.1±0.06 | 0.06±0.01 | 0.03±0.01 |
| *Pedobacter* | 0.07±0.02 | 0.07±0.03 | 0.06±0.01 | 0.06±0.01 | 0.05±0.02 |
| *Myceligenerans* | | 0.01±0.01 | | 0.14±0.17 | 0.01±0.02 |
| *Gordonia* | | | | 0.09±0.17 | 0.07±0.13 |
| *Promicromonospora* | | | 0.02±0.02 | 0.1±0.12 | 0.04±0.03 |
| *Microbacterium* | 0.04±0.01 | 0.01±0.01 | 0.02±0.02 | 0.08±0.09 | 0.03±0.04 |
| *Buttiauxella* | 0.01±0 | 0.03±0 | 0.04±0.02 | 0.03±0.01 | 0.04±0.01 |
| *Isoptericola* | | | | 0.07±0.12 | 0.06±0.03 |
| *Brevibacterium* | 0.01±0 | | | 0.11±0.21 | |
| *Pseudonocardia* | | | | | 0.08±0.04 |
| *Agromyces* | 0.02±0.01 | | 0.06±0.03 | 0.01±0.01 | 0.01±0.01 |
| *Variovorax* | 0.02±0.01 | 0.03±0.02 | 0.02±0.01 | 0.02±0.01 | |
| *Skermanella* | | | 0.06±0.1 | | |
| *Caenimicrobium* | | | 0.06±0.06 | | |
| *Saccharomonospora* | | | | | 0.05±0.09 |
| *Phyllobacterium* | 0.01±0.01 | 0.01±0 | 0.02±0.01 | 0.02±0.01 | 0.01±0 |
| *Stenotrophomonas* | 0.01±0 | 0.02±0.01 | 0.01±0.01 | 0.01±0.01 | 0.01±0.01 |
| *Staphylococcus* | | | | 0.04±0.08 | |
| *Kangiella* | 5.41±3.07 | 0.02±0.01 | 0.01±0 | 0.01±0.01 | 0.01±0.01 |
| Saccharibacteria g i s | 2.33±1.4 | 0.01±0.01 | 0.01±0.01 | 0.01±0.01 | 0.01±0 |
| *Pseudidiomarina* | 2.6±0.54 | 0.01±0 | 0.01±0.01 | 0.01±0.01 | 0.01±0.01 |
| *Halomonas* | 0.86±0.11 | | 0.01±0.01 | 0.01±0.01 | 0.01±0.01 |
| Others | 8.14±1.16 | 0.09±0.03 | 0.15±0.04 | 0.09±0.04 | 0.09±0.03 |
| Unclassified | 77.9±2.6 | 18.93±5.83 | 12.78±1 | 5.06±1.93 | 4.9±1.65 |

Table S6. Total, rare, and unique OTUs obtained through the culture-dependent approach using TSA (tryptic soy agar), SEM (soil extract medium), R2A (Reasoner's 2A), and 1:20 R2A (Reasoner's 2A diluted 20-fold) from agricultural, forest, and contaminated soils. Values represent mean ± standard deviation (n = 4) and, for each soil, different letters indicate significant differences ($P < 0.05$) according to ANOVA followed by Tukey's HSD test. Values represent means ± standard deviations.

| Medium | Total OTUs | Rare OTUs | Unique OTUs |
|---|---|---|---|
| Agricultural soil | | | |
| TSA | 120.75 ± 3.95 a | 23.00 ± 2.94 a | 52.25 ± 6.70 a |
| SEM | 251 ± 7.70 c | 38.75 ± 6.65 b | 85.75 ± 8.62 b |
| R2A | 160.50 ± 4.04 b | 27.75 ± 5.74 ab | 54.75 ± 10.90 a |
| 1:20 R2A | 179.25 ± 18.95 b | 32.00 ± 6.27 ab | 61.75 ± 3.30 a |
| Forest soil | | | |
| TSA | 87.00 ± 6.06 b | 19.50 ± 1.29 ab | 54.25 ± 6.60 a |
| SEM | 145.75 ± 26.22 c | 37.25 ± 3.95 c | 134.00 ± 12.20 c |
| R2A | 55.75 ± 6.95 a | 13.25 ± 3.40 a | 99.75 ± 13.65 b |
| 1:20 R2A | 80.75 ± 4.03 ab | 22.00 ± 2.94 b | 113.25 ± 16.64 b |
| Contaminated soil | | | |
| TSA | 65.67 ± 7.64 a | 16.33 ± 5.86 a | 50.67 ± 2.31 a |
| SEM | 107.00 ± 5.89 b | 26.00 ± 3.83 b | 79.00 ± 8.04 c |
| R2A | 73.75 ± 4.57 a | 17.25 ± 4.57 ab | 60.25 ± 4.50 b |
| 1:20 R2A | 74.75 ± 7.63 a | 18.00 ± 1.83 ab | 64.50 ± 5.74 b |

Table S7. Taxonomic composition at genus level of the rare bacterial fraction isolated using TSA (tryptic soy agar), SEM (soil extract medium), R2A (Reasoner's 2A), and 1:20 R2A (Reasoner's 2A diluted 20-fold) from agricultural soil. Values represent percentages.

| Genus | TSA | SEM | R2A | 1:20 R2A |
|---|---|---|---|---|
| *Bacillus* | 22.30 | 2.06 | 28.70 | 21.30 |
| *Nocardioides* | 0.12 | 46.20 | 2.99 | 12.10 |
| *Xanthomonas* | 0.55 | 5.47 | 1.48 | 7.64 |
| *Microbacterium* | 3.99 | 2.08 | 4.34 | 4.16 |
| *Chitinophaga* | 0.07 | 8.99 | 0.05 | 4.70 |
| *Pedobacter* | 0.07 | 0.14 | 12.70 | 0.20 |
| *Paenibacillus* | 7.47 | 0.63 | 0.36 | 2.69 |
| *Solibacillus* | 10.80 | 0.11 | 0.08 | 0.15 |
| *Acinetobacter* | 5.11 | 0.52 | 1.86 | 1.91 |
| *Lysinibacillus* | 6.62 | | 0.05 | |
| *Williamsia* | | 0.14 | 0.10 | 5.97 |
| *Pseudoxanthomonas* | 0.12 | 4.79 | 0.31 | 0.93 |
| *Delftia* | | | 1.91 | 4.02 |
| *Stenotrophomonas* | 0.99 | 2.60 | 0.89 | 1.22 |
| *Agromyces* | 0.10 | 0.75 | 3.04 | 1.76 |
| *Advenella* | 0.87 | 1.33 | 2.14 | 1.18 |
| *Sphingobacterium* | 3.85 | 0.11 | 0.95 | 0.15 |
| *Rhizobium* | 3.03 | 0.27 | 1.17 | 0.10 |
| *Roseimicrobium* | 0.02 | 2.96 | 0.03 | |
| *Flavobacterium* | 0.25 | 1.40 | 0.89 | 0.44 |
| *Cupriavidus* | 0.15 | 0.52 | 0.64 | 1.32 |
| *Curtobacterium* | 1.07 | 0.25 | 0.26 | 0.74 |
| *Streptomyces* | 0.37 | 0.84 | 0.15 | 0.69 |
| *Sporosarcina* | 0.10 | 0.20 | 0.66 | 0.34 |
| *Fictibacillus* | | 0.50 | | 0.44 |
| *Mesobacillus* | 0.02 | 0.36 | | 0.54 |
| *Chryseobacterium* | | | | 0.88 |
| *Cellulomonas* | 0.20 | 0.25 | 0.03 | 0.15 |
| *Plantibacter* | 0.30 | 0.02 | 0.10 | 0.20 |
| *Luteibacter* | 0.02 | | | 0.49 |
| *Flavisolibacter* | | 0.50 | | |
| *Bhargavaea* | 0.50 | | | |
| *Knoellia* | | 0.23 | 0.15 | |
| *Neobacillus* | | 0.14 | 0.03 | 0.20 |
| Spartobacteria g i s | | 0.20 | | 0.10 |
| *Cellulosimicrobium* | 0.07 | 0.02 | 0.05 | 0.10 |
| *Aeromicrobium* | | 0.11 | 0.03 | 0.10 |
| *Staphylococcus* | 0.02 | 0.11 | | 0.10 |
| *Oceanobacillus* | 0.05 | 0.05 | 0.05 | 0.05 |
| *Oerskovia* | 0.12 | 0.05 | 0.03 | |
| *Gp6* | 0.02 | 0.07 | | 0.10 |
| *Solirubrobacter* | | 0.09 | 0.03 | 0.05 |
| *Niabella* | | 0.14 | 0.03 | |
| *Gp4* | | 0.05 | | 0.10 |
| *Leifsonia* | | 0.05 | | 0.10 |
| Subdivision3 g i s | | 0.05 | | 0.10 |
| *Glycomyces* | | 0.09 | | 0.05 |
| *Alkalihalobacillus* | 0.12 | | | |
| *Frigoribacterium* | 0.12 | | | |
| *Labedella* | | 0.02 | 0.05 | 0.05 |
| *Caballeronia* | | 0.09 | 0.03 | |
| *Nocardiopsis* | | 0.09 | 0.03 | |
| *Pseudomonas* | | 0.09 | 0.03 | |
| *Chryseolinea* | 0.02 | | 0.03 | 0.05 |
| *Thermomonas* | | | | 0.10 |
| WPS-1 g i s | | | | 0.10 |
| *Paraburkholderia* | | | 0.03 | 0.05 |

| | | | | |
|---|---|---|---|---|
| *Agrococcus* | | 0.07 | | |
| *Singulisphaera* | | 0.07 | | |
| *Hymenobacter* | | | | 0.05 |
| *Edaphobacter* | | 0.02 | 0.03 | |
| *Schlesneria* | | 0.05 | | |
| *Arthrobacter* | | | 0.03 | |
| Gp17 | | | 0.03 | |
| *Streptococcus* | | | 0.03 | |
| *Lacipirellula* | 0.02 | | | |
| *Leucobacter* | 0.02 | | | |
| *Ferruginibacter* | | 0.02 | | |
| *Gemmata* | | 0.02 | | |
| *Larkinella* | | 0.02 | | |
| *Mycobacterium* | | 0.02 | | |
| *Saccharopolyspora* | | 0.02 | | |
| *Terrimonas* | | 0.02 | | |
| *Verrucomicrobium* | | 0.02 | | |
| Unclassified | 30.40 | 13.90 | 33.40 | 22.10 |

Table S8. Taxonomic composition at genus level of the rare bacterial fraction isolated using TSA (tryptic soy agar), SEM (soil extract medium), R2A (Reasoner's 2A), and 1:20 R2A (Reasoner's 2A diluted 20-fold) from forest soil. Values represent percentages.

| Genus | TSA | SEM | R2A | 1:20 R2A |
|---|---|---|---|---|
| *Pedobacter* | 0.11 | 33.30 | 37.30 | 44.80 |
| *Phyllobacterium* | 0.11 | 22.00 | 15.60 | 5.58 |
| *Chitinophaga* | 0.13 | 11.80 | 0.98 | 25.70 |
| *Sphingobacterium* | 34.00 | 0.14 | 1.09 | |
| *Sporosarcina* | 20.30 | 0.05 | | |
| *Pseudomonas* | 3.74 | 6.89 | 1.41 | 7.76 |
| *Oceanobacillus* | 8.13 | | | |
| *Agromyces* | 0.53 | 0.38 | 5.64 | 0.51 |
| *Flavobacterium* | 0.06 | 2.36 | 3.28 | 0.91 |
| *Staphylococcus* | 5.10 | 0.02 | 0.08 | 0.05 |
| *Nocardioides* | 0.47 | 2.85 | 0.85 | 0.84 |
| *Streptomyces* | 0.21 | 1.33 | 1.34 | 1.83 |
| *Chryseobacterium* | 3.80 | 0.14 | 0.37 | 0.24 |
| *Neobacillus* | 0.11 | 2.17 | 0.04 | 0.01 |
| *Mesobacillus* | | 0.53 | 1.02 | 0.69 |
| *Paraburkholderia* | 0.11 | 1.06 | 0.07 | 0.99 |
| *Lysobacter* | 0.09 | 1.11 | 0.48 | 0.24 |
| *Dyadobacter* | 0.02 | 1.25 | 0.07 | 0.25 |
| *Leifsonia* | 0.06 | 0.18 | 0.78 | 0.19 |
| *Paenibacillus* | 0.09 | 0.74 | 0.33 | 0.03 |
| *Nocardia* | | 0.73 | 0.01 | 0.08 |
| *Bacillus* | 0.17 | 0.12 | 0.38 | 0.15 |
| *Caballeronia* | | 0.35 | | 0.43 |
| *Mucilaginibacter* | | 0.62 | | 0.10 |
| *Mycobacterium* | 0.04 | 0.09 | 0.07 | 0.33 |
| *Roseimicrobium* | 0.02 | 0.45 | | |
| *Pseudoxanthomonas* | 0.15 | 0.13 | 0.03 | 0.04 |
| *Kitasatospora* | 0.02 | 0.02 | 0.08 | 0.20 |
| *Microbacterium* | 0.15 | 0.05 | 0.08 | 0.03 |
| *Mesorhizobium* | | 0.17 | 0.01 | 0.11 |
| *Terriglobus* | | 0.22 | | 0.04 |
| *Luteibacter* | 0.17 | 0.07 | 0.01 | |
| *Singulisphaera* | | 0.24 | | 0.01 |
| *Terrabacter* | | 0.09 | 0.04 | 0.06 |
| *Cellulomonas* | | 0.02 | 0.07 | 0.10 |
| *Solibacillus* | 0.04 | 0.04 | 0.07 | 0.04 |
| *Microlunatus* | | 0.09 | 0.08 | 0.01 |
| *Williamsia* | 0.02 | 0.01 | 0.06 | 0.05 |
| *Oerskovia* | 0.13 | | | |
| *Pantoea* | 0.13 | | | |
| Spartobacteria g i s | 0.09 | 0.04 | | |
| *Embleya* | | 0.04 | 0.06 | 0.01 |
| *Solirubrobacter* | 0.02 | 0.04 | | 0.03 |
| *Olivibacter* | 0.04 | 0.01 | | |
| *Promicromonospora* | 0.04 | 0.01 | | |
| *Stenotrophomonas* | 0.04 | 0.01 | | |
| *Acidovorax* | | 0.01 | 0.04 | |
| *Massilia* | | 0.05 | | |
| *Niastella* | | 0.05 | | |
| *Pandoraea* | 0.04 | | | |
| *Skermanella* | 0.04 | | | |
| *Gemmata* | | 0.04 | | |
| *Taibaiella* | | 0.04 | | |
| *Acinetobacter* | | 0.02 | | |
| *Nonomuraea* | | 0.01 | | 0.01 |
| Gp4 | 0.02 | | | |

| | | | | |
|---|---|---|---|---|
| *Kribbella* | 0.02 | | | |
| *Peribacillus* | 0.02 | | | |
| *Tsukamurella* | | | | 0.02 |
| *Daejeonella* | | 0.01 | | |
| *Nitrospira* | | 0.01 | | |
| *Stenotrophobacter* | | 0.01 | | |
| *Pseudonocardia* | | | | 0.01 |
| Unclassified | 21.50 | 7.81 | 28.10 | 7.56 |

Table S9. Taxonomic composition at genus level of the rare bacterial fraction isolated using TSA (tryptic soy agar), SEM (soil extract medium), R2A (Reasoner's 2A), and 1:20 R2A (Reasoner's 2A diluted 20-fold) from contaminated soil. Values represent percentages.

| Genus | TSA | SEM | R2A | 1:20 R2A |
|---|---|---|---|---|
| *Bacillus* | 33.00 | 0.60 | 34.20 | 14.30 |
| *Solibacillus* | 37.00 | 0.22 | 0.05 | 0.17 |
| *Streptomyces* | 1.88 | 3.37 | 14.00 | 14.70 |
| *Fictibacillus* | 0.37 | 0.33 | 23.10 | 9.87 |
| *Janibacter* | 4.15 | 12.00 | 2.05 | 0.07 |
| *Neobacillus* | 0.14 | 1.14 | 1.92 | 14.70 |
| *Virgibacillus* | 9.35 | | 0.05 | 0.02 |
| *Georgenia* | 0.03 | 7.72 | 0.38 | 0.55 |
| *Alkalihalobacillus* | 0.06 | 1.68 | 1.31 | 5.13 |
| *Caenimicrobium* | | 2.17 | | |
| *Isoptericola* | | 0.05 | 0.80 | 1.02 |
| *Flavobacterium* | 0.46 | 0.76 | 0.49 | 0.13 |
| *Pseudomonas* | 0.48 | 0.60 | 0.29 | 0.37 |
| *Pseudonocardia* | | 0.11 | | 1.40 |
| *Brevibacterium* | | 0.05 | 1.40 | 0.02 |
| *Gordonia* | 0.03 | | 1.09 | |
| *Stenotrophomonas* | 0.26 | 0.44 | 0.07 | 0.20 |
| *Luteimonas* | | 0.76 | 0.05 | 0.02 |
| *Nocardioides* | 0.09 | 0.27 | 0.11 | 0.17 |
| *Agromyces* | 0.03 | 0.27 | 0.11 | 0.15 |
| *Chitinophaga* | 0.06 | 0.27 | 0.09 | |
| *Serinicoccus* | | | 0.15 | 0.22 |
| *Aeromonas* | 0.09 | 0.16 | 0.04 | 0.05 |
| *Peribacillus* | 0.09 | 0.05 | 0.04 | 0.07 |
| *Pedobacter* | 0.03 | 0.11 | 0.04 | 0.02 |
| *Blastopirellula* | 0.03 | 0.16 | | |
| *Clostridium* s e | 0.11 | | | |
| *Microbulbifer* | | 0.11 | | |
| *Mycobacterium* | | 0.05 | | 0.02 |
| Spartobacteria g i s | | 0.05 | | 0.02 |
| *Anseongella* | | 0.05 | | |
| *Marinobacter* | | 0.05 | | |
| *Saccharomonospora* | | | | 0.05 |
| *Solirubrobacter* | | | | 0.05 |
| *Paraburkholderia* | | | | 0.02 |
| *Variovorax* | | | | 0.02 |
| Unclassified | 12.30 | 66.40 | 18.20 | 36.40 |

Table S10. Taxonomic composition at genus level of the unique bacterial fraction isolated using TSA (tryptic soy agar), SEM (soil extract medium), R2A (Reasoner's 2A), and 1:20 R2A (Reasoner's 2A diluted 20-fold) from agricultural soil. Values represent percentages.

| Genus | TSA | SEM | R2A | 1:20 R2A |
|---|---|---|---|---|
| *Bacillus* | 68.40 | 3.02 | 45.70 | 8.72 |
| *Staphylococcus* | 0.35 | 0.32 | 0.36 | 39.30 |
| *Olivibacter* | 0.14 | 23.30 | 0.53 | |
| *Microbacterium* | 0.28 | 1.62 | 4.08 | 15.80 |
| *Paenibacillus* | 5.28 | 3.13 | 6.91 | 2.31 |
| *Nocardioides* | | 4.75 | 0.18 | 1.78 |
| *Pseudomonas* | 0.90 | 1.30 | 3.19 | 0.89 |
| *Fictibacillus* | | 5.72 | 0.18 | |
| *Devosia* | 0.07 | 1.08 | 1.24 | 2.31 |
| *Brucella* | 3.75 | 0.54 | | |
| *Sphingobacterium* | 2.50 | 0.76 | 0.71 | |
| *Aeromicrobium* | | 0.32 | 3.37 | |
| *Stenotrophomonas* | 0.63 | 0.43 | 1.42 | 1.07 |
| *Chitinophaga* | 0.21 | 0.65 | 0.53 | 0.71 |
| *Flavobacterium* | 0.69 | 0.43 | 0.18 | 0.71 |
| *Agrobacterium* | 0.35 | 0.11 | 0.53 | 0.53 |
| *Lysobacter* | | 1.51 | | |
| Spartobacteria_g i s | 0.07 | 0.65 | | 0.53 |
| *Mucilaginibacter* | 0.07 | 1.08 | | |
| *Microlunatus* | 0.07 | | 0.71 | |
| *Pedobacter* | | 0.76 | | |
| Gp6 | 0.07 | 0.32 | | 0.36 |
| *Brevibacillus* | | 0.22 | 0.53 | |
| *Sphingobium* | | | 0.71 | |
| *Dyella* | | 0.32 | 0.36 | |
| Subdivision3 g i s | | 0.32 | 0.18 | 0.18 |
| *Arthrobacter* | | 0.11 | 0.18 | 0.36 |
| *Streptomyces* | 0.21 | 0.11 | | 0.18 |
| *Cutibacterium* | 0.07 | 0.11 | | 0.18 |
| *Sporosarcina* | 0.07 | | | 0.18 |
| *Mesorhizobium* | | | | 0.18 |
| *Ferruginibacter* | 0.07 | 0.11 | | |
| *Latescibacteria* g i s | | | 0.18 | |
| *Rhizobium* | | | 0.18 | |
| *Edaphobacter* | | 0.11 | | |
| *Nemorincola* | | 0.11 | | |
| *Streptophyta* | | 0.11 | | |
| *Yinghuangia* | | 0.11 | | |
| Gp5 | 0.07 | | | |
| Unclassified | 15.70 | 46.40 | 27.80 | 23.70 |

Table S11. Taxonomic composition at genus level of the unique bacterial fraction isolated using TSA (tryptic soy agar), SEM (soil extract medium), R2A (Reasoner's 2A), and 1:20 R2A (Reasoner's 2A diluted 20-fold) from forest soil. Values represent percentages.

| Genus | TSA | SEM | R2A | 1:20 R2A |
|---|---|---|---|---|
| *Bacillus* | 18.20 | 35.40 | 4.12 | 3.67 |
| *Pseudomonas* | 15.90 | 7.73 | 20.50 | 20.10 |
| *Paenibacillus* | 7.65 | 2.77 | 6.72 | 0.25 |
| *Lysobacter* | 0.38 | 2.46 | 6.84 | 5.76 |
| *Flavobacterium* | 0.38 | 3.35 | 2.41 | 7.09 |
| *Sporosarcina* | 8.22 | 0.04 | | |
| *Mucilaginibacter* | 0.38 | 1.79 | 0.19 | 4.18 |
| *Fictibacillus* | 0.38 | 0.72 | 2.60 | 1.77 |
| *Pedobacter* | 0.38 | 1.38 | 2.34 | 1.33 |
| *Acinetobacter* | 2.68 | 0.27 | 1.14 | 0.25 |
| *Rhizobium* | 0.57 | 0.45 | 2.53 | 0.70 |
| *Arthrobacter* | 3.63 | 0.04 | 0.19 | |
| *Polaromonas* | | 0.67 | 0.95 | 1.14 |
| *Brucella* | | 0.31 | 0.76 | 1.27 |
| *Luteibacter* | | 1.07 | 0.13 | 1.08 |
| *Chryseobacterium* | 1.34 | 0.40 | 0.19 | 0.19 |
| *Nocardioides* | 0.96 | 0.45 | 0.25 | 0.38 |
| *Xanthomonas* | 0.96 | 0.04 | | 0.95 |
| *Streptomyces* | 0.19 | 0.54 | 0.32 | 0.63 |
| *Kangiella* | | 0.27 | 0.63 | 0.76 |
| *Pseudidiomarina* | | 0.09 | 0.63 | 0.51 |
| *Dyadobacter* | | 0.67 | 0.32 | 0.13 |
| *Agromyces* | 0.57 | 0.09 | 0.06 | 0.32 |
| *Saccharibacteria gis* | | 0.18 | 0.44 | 0.38 |
| *Phyllobacterium* | | 0.45 | | 0.44 |
| *Stenotrophomonas* | 0.77 | 0.04 | | |
| *Marmoricola* | | 0.31 | 0.06 | 0.38 |
| *Filimonas* | | 0.27 | | 0.38 |
| *Spartobacteria gis* | | 0.63 | | |
| *Gracilimonas* | | 0.13 | 0.25 | 0.19 |
| Clostridium_sensu_stricto | 0.57 | | | |
| *Microbacterium* | 0.38 | 0.04 | 0.06 | 0.06 |
| *Skermanella* | 0.38 | 0.09 | | |
| *Streptococcus* | 0.38 | | 0.06 | |
| *Taibaiella* | | 0.31 | 0.06 | 0.06 |
| *Chitinophaga* | 0.38 | 0.04 | | |
| *Massilia* | 0.38 | 0.04 | | |
| *Agrobacterium* | 0.38 | | | |
| *Halomonas* | | | 0.13 | 0.25 |
| *Comamonas* | 0.19 | 0.04 | 0.13 | |
| *Ornithinimicrobium* | | 0.04 | 0.13 | 0.19 |
| *Marinobacter* | | 0.04 | 0.06 | 0.25 |
| *Nocardia* | | 0.36 | | |
| *Cutibacterium* | 0.19 | 0.04 | | 0.06 |
| *Janibacter* | | 0.04 | 0.13 | 0.13 |
| *Delftia* | 0.19 | | | 0.06 |
| *Mycobacterium* | | | | 0.25 |
| *Aeromicrobium* | 0.19 | 0.04 | | |
| *Cellvibrio* | 0.19 | 0.04 | | |
| *Cystobacter* | 0.19 | 0.04 | | |
| *Pseudoxanthomonas* | 0.19 | 0.04 | | |
| *Rhodanobacter* | | 0.04 | | 0.19 |
| *Sphingobium* | | 0.22 | | |
| Gp1 | 0.19 | | | |
| *Terrimicrobium* | 0.19 | | | |
| *Egicoccus* | | | 0.06 | 0.13 |

| | | | | |
|---|---|---|---|---|
| *Pseudactinotalea* | | | 0.13 | 0.06 |
| *Cyclobacterium* | | | 0.19 | |
| *Virgibacillus* | | 0.04 | 0.06 | 0.06 |
| *Pelagibius* | | 0.13 | | |
| *Spirosoma* | | 0.13 | | |
| *Steroidobacter* | | 0.13 | | |
| *Dietzia* | | | | 0.13 |
| *Gordonia* | | | | 0.13 |
| *Alkalihalobacillus* | | | 0.06 | 0.06 |
| *Curtobacterium* | | | 0.06 | 0.06 |
| *Gimesia* | | | 0.06 | 0.06 |
| *Planktosalinus* | | | 0.06 | 0.06 |
| *Pusillimonas* | | | 0.06 | 0.06 |
| *Tumebacillus* | | 0.04 | | 0.06 |
| *Flavisolibacter* | | 0.09 | | |
| *Lysinibacillus* | | 0.09 | | |
| *Alcanivorax* | | | 0.06 | |
| *Arenibacter* | | | 0.06 | |
| *Brevibacterium* | | | 0.06 | |
| *Kocuria* | | | 0.06 | |
| *Listeria* | | | 0.06 | |
| *Salinimicrobium* | | | 0.06 | |
| *Thiohalobacter* | | | 0.06 | |
| *Aequorivita* | | | | 0.06 |
| *Cellulosimicrobium* | | | | 0.06 |
| *Edaphobacter* | | | | 0.06 |
| *Georgenia* | | | | 0.06 |
| *Methyloceanibacter* | | | | 0.06 |
| *Muricauda* | | | | 0.06 |
| *Rubinisphaera* | | | | 0.06 |
| *Salegentibacter* | | | | 0.06 |
| *Sediminibacter* | | | | 0.06 |
| *Solibacillus* | | | | 0.06 |
| *Thioalkalivibrio* | | | | 0.06 |
| *Advenella* | | 0.04 | | |
| *Chryseolinea* | | 0.04 | | |
| *Geobacillus* | | 0.04 | | |
| Gp6 | | 0.04 | | |
| *Methylophaga* | | 0.04 | | |
| *Methylotenera* | | 0.04 | | |
| *Microbulbifer* | | 0.04 | | |
| *Planococcus* | | 0.04 | | |
| *Rhodopirellula* | | 0.04 | | |
| *Segetibacter* | | 0.04 | | |
| *Streptosporangium* | | 0.04 | | |
| Unclassified | 31.90 | 34.30 | 43.40 | 42.70 |

Table S12. Taxonomic composition at genus level of the unique bacterial fraction isolated using TSA (tryptic soy agar), SEM (soil extract medium), R2A (Reasoner's 2A), and 1:20 R2A (Reasoner's 2A diluted 20-fold) from contaminated soil. Values represent percentages.

| Genus | TSA | SEM | R2A | 1:20 R2A |
|---|---|---|---|---|
| *Bacillus* | 33.90 | 0.20 | 16.80 | 2.46 |
| *Streptomyces* | 1.76 | 4.76 | 7.29 | 23.10 |
| *Lysobacter* | 25.20 | 4.26 | 0.12 | 0.32 |
| *Paenibacillus* | 0.62 | 0.51 | 24.20 | 1.19 |
| *Pseudactinotalea* | | 11.10 | 2.53 | 5.87 |
| *Dietzia* | 1.87 | 14.00 | 0.06 | 1.98 |
| *Solibacillus* | 14.50 | | | 0.16 |
| *Pseudomonas* | 0.52 | 12.70 | 0.63 | 0.79 |
| *Cellulosimicrobium* | 0.31 | | 0.12 | 16.90 |
| *Tsukamurella* | | | 11.00 | 0.08 |
| *Fictibacillus* | | 0.10 | 0.69 | 10.05 |
| *Promicromonospora* | | 1.11 | 3.73 | 1.98 |
| *Myceligenerans* | 0.31 | 0.10 | 5.40 | 0.79 |
| *Microbacterium* | 0.31 | 1.22 | 3.27 | 1.75 |
| *Peribacillus* | 0.21 | | 4.19 | |
| *Skermanella* | | 4.26 | 0.06 | 0.08 |
| *Gordonia* | 0.10 | | | 3.49 |
| *Saccharomonospora* | | 0.30 | | 2.46 |
| *Staphylococcus* | | 0.20 | 1.61 | |
| *Nocardioides* | | 0.10 | 0.12 | 2.51 |
| *Agrococcus* | 0.52 | 0.20 | 0.06 | 0.32 |
| *Flavobacterium* | 0.10 | 0.61 | 0.17 | 0.16 |
| *Mesobacillus* | 1.04 | | | |
| *Oceanobacillus* | 1.04 | | | |
| *Lysinibacillus* | | | 0.80 | |
| *Neobacillus* | | | | 0.71 |
| *Micrococcus* | | 0.61 | | |
| *Georgenia* | 0.21 | | | 0.32 |
| *Metabacillus* | | | 0.52 | |
| *Rhizobium* | | 0.41 | | 0.08 |
| *Pedobacter* | 0.10 | 0.10 | 0.06 | 0.16 |
| *Janibacter* | | 0.41 | | |
| *Amycolatopsis* | | | | 0.40 |
| *Microlunatus* | | | 0.17 | 0.16 |
| *Martelella* | | 0.30 | | |
| *Rhodococcus* | 0.21 | | | 0.08 |
| *Agromyces* | 0.10 | | 0.17 | |
| *Stenotrophomonas* | | 0.20 | 0.06 | |
| *Chitinophaga* | 0.10 | | 0.06 | 0.08 |
| *Streptosporangium* | | | | 0.24 |
| *Leifsonia* | | | 0.06 | 0.16 |
| *Luteibacter* | | 0.20 | | |
| *Nocardiopsis* | | 0.20 | | |
| *Schlesneria* | | 0.20 | | |
| *Sphingobacterium* | 0.10 | | | 0.08 |
| *Ureibacillus* | 0.10 | | | 0.08 |
| *Arthrobacter* | | | | 0.16 |
| *Marmoricola* | | | | 0.16 |
| *Kitasatospora* | | | 0.06 | 0.08 |
| *Acinetobacter* | 0.10 | | | |
| *Nocardia* | 0.10 | | | |
| *Aeromicrobium* | | 0.10 | | |
| *Mesorhizobium* | | 0.10 | | |
| *Psychrobacillus* | | 0.10 | | |
| Spartobacteria g i s | | 0.10 | | |
| *Mycobacterium* | | | | 0.08 |

| | | | | |
|---|---|---|---|---|
| *Ramlibacter* | | | | 0.08 |
| *Tumebacillus* | | | | 0.08 |
| *Williamsia* | | | | 0.08 |
| *Yinghuangia* | | | | 0.08 |
| *Curtobacterium* | | | 0.06 | |
| *Terrabacter* | | | 0.06 | |
| Unclassified | 16.60 | 41.20 | 15.80 | 20.20 |
